# Genomic prediction of agronomic traits in perennial ryegrass (*Lolium perenne* L.) and genotype x environment interactions at the limit of the species distribution

**DOI:** 10.1101/2025.03.18.642199

**Authors:** Natasha H. Johansen, Andrea Bellucci, Pernille B. Hansen, Petter Marum, Helga Amdahl, Kristin Håland Gylstrøm, Odd Arne Rognli, Vilma Kemešytė, Gintaras Brazauskas, Morten Greve, Christer Persson, Mika Isolahti, Áslaug Helgadóttir, Rene Aavola, Torben Asp, Guillaume P. Ramstein

## Abstract

**Background:** In breeding the aim is to identify and accumulate beneficial variants. However, detection of these variants may be challenging in the presence of extensive genotype x environment interactions (GxE), as variant effects will be conditional on environment.

**Methods:** The study assesses the performance of **264** diploid perennial ryegrass accessions in a multi-environment field trial. We investigate the extent of GxE, for yield (total dry matter) and persistence traits, i.e. winter kill and spring cover, under environmental conditions experienced in Nordic and Baltic regions at the limit of the species distribution. Two different approaches to modelling GxE were tested: reaction norm and envirotyping, and the models were validated under three different breeding scenarios.

**Results:** Our analysis documented the presence of significant GxE interaction for all traits investigated in the study. Validation showed improvements in prediction accuracy when accounting for GxE: up to 4% for yield when predicting in unobserved environments, and up to 22% and 9% for spring cover and winter kill, respectively when predicting unobserved germplasm. Genome-wide-association-studies (GWAS) were utilized to detect genetic variants with marginal effects (environment-independent effect) and conditional effects (environment-dependent effects). Results showed the presence of large-effect genetic variants with marginal effects, in addition to few QTL whose effects were adaptive under specific environmental conditions while neutral or deleterious under different environmental conditions.

**Conclusion:** This study demonstrates the usefulness and limitations of genomic prediction models for predicting GxE in highly diverse samples, and describes the extent of GxE interactions at the limit of species distribution for perennial ryegrass. Finally, our study points toward adaptive variation, which may enhance persistence of perennial ryegrass populations in Nordic and Baltic growing conditions.

## 1. Introduction

Climate changes will in the future lead to increased fluctuations in temperature and rainfall, and therefore, more unpredictable weather patterns, which may challenge plant breeders in the future (Raza et al. 2019). Consequently, to ensure that future food demand can be supported, it’s necessary to adapt current food production practices and develop high yielding crop varieties that are adapted to unpredictable weather patterns (Kahiluoto et al. 2019). Different selection strategies have been developed to facilitate efficient evaluation of superior breeding material. These selection strategies include, among other, Genomic Prediction (GP) (Meuwissen et al. 2001) which has become an important tool to predict the genetic performance, i.e. breeding values, of breeding material from genetic markers. Plant breeders evaluate candidate lines/populations in multiple environmental trials (MET) to ensure that these lines/populations have stable performance, i.e., broad adaptation, across a range of different environments. These environments are referred to as the target population of environments (TPE), which encompasses the intended production environments of the lines. However, broad adaptation does not necessarily imply superior performance across environments, as some lines/populations may express superior performance in a narrow range of environments, due to local adaptation to specific environmental conditions (Ågren and Schemske 2012; Ågren et al. 2013; Lasky et al. 2017; Blanco-Pastor et al. 2021). Hence, differences in phenotypic performance between lines/populations are not exclusively caused by genetic effects but are also affected by genotype by environment interactions (GxE), i.e., differential responses of genotypes over an environmental gradient. GxE can be expressed as (i) non-cross-over interaction, i.e., genotypes maintain their rankings, and thus undergoes scale changes. (ii) cross-over interaction where the rankings of the genotypes are changed, which occur when genotype rankings are environment-dependent (El-Soda et al. 2014). Traditionally, plant breeders have ignored or minimised GxE interactions to avoid the challenge of modelling this component, however studies show that the prediction accuracy of GP models can be enhanced in MET, by accounting for GxE (Jarquín et al. 2014, 2017; Cuevas et al. 2017; Millet et al. 2019; Jarquin et al. 2020). Furthermore, integration of GxE in GP models allows for prediction of phenotypic performance in unobserved environments, which can be of great advantage to breeders. Different approaches have been developed to integrate GxE into genomic prediction models. These approaches include, amongst others extensions of Genomic Best Linear Unbiased Prediction models (GBLUP) (Habier et al. 2007), which accounts for the covariance of environmental values and GxE interactions: reaction norm (Jarquín et al. 2014) and envirotyping approaches (Xu 2016; Costa-Neto et al. 2021cb, a). These approaches differ regarding how the environmental covariates are transformed. The reaction norm approach models GxE by genotypes’ linear response to environmental covariates.

Thus, in reaction-norm models, more complex interactions, i.e., non-linear interactions, are not captured (Jarquín et al. 2014). Envirotyping is an approach by which environments are characterised by determining the environmental covariates that affect a given plant species’ growth (Xu 2016; Costa-Neto et al. 2021cb, a). The envirotyping approach is comparable to the reaction norm approach but differs in how environmental relatedness is calculated. For the reaction norm approach environmental relatedness is based on quantitative descriptors (e.g., mean temperature) whereas for the envirotyping approach environmental gradients are partitioned into environmental zones, or typologies, according to physiological expectations about survival, tolerance, or optimal growth for the given crop species, as inspired by Shelford’s Law of Tolerance (Shelford 1931). The number of occurrences of these environmental zones are determined for each environment, which allows for environmental relatedness to be based on the frequency of each environmental zone that an environment experiences and therefore also informs about the value of the environment for the given crop (Costa-Neto et al. 2021c, b, a). The aim of GP is to identify superior lines based on genome-wide variants; however, GP cannot be used to detect impactful variants at a resolution that allows for detection of potential QTL (Quantitative Trait Loci). Genome-Wide-Association-studies (GWAS) are commonly used for this instead (Liu and Yan 2019; Tibbs Cortes et al. 2021). Linkage disequilibrium (LD), i.e., non-random association of alleles at different loci, is integral for GWAS as the power and ability to detect genotype-phenotype associations is lower when neutral alleles are not associated with causal variants (Flint-Garcia et al. 2003). Association analysis in populations with rapid LD decay therefore requires high marker densities, as genetic variants will be linked together in smaller blocks, compared to populations with more extensive LD. GWAS are commonly used to detect QTL based on their average association with traits, also termed marginal effects. However, GWAS are also a useful tool for the detection of variants with conditional effects, which can occur due to local adaptation. These may be detected based on genome-environment associations (Lasky et al. 2017; Li et al. 2021). Conditional neutrality of variant effects and genetic trades-offs (maladaptive pleiotropy) may both produce conditional variant effects, where conditional neutrality implies that a variant may be beneficial or deleterious under certain environmental conditions, while effectively neutral in others, while alleles showing maladaptive pleiotropy will confer a fitness advantage under some environmental conditions and a fitness disadvantage in others (Hereford 2009; Anderson et al. 2013). Perennial ryegrass (*Lolium perenne* L.) is an economically important forage grass used for animal feed in dairy production, where it is valued because of its high yields and nutritional value. Additionally, the crop expresses a high level of genetic diversity, due to its obligate outbreeding nature (Brazauskas et al. 2011). Nordic regions, which have previously been unsuitable for production of perennial ryegrass, can become fit for production of certain robust and cold-tolerant varieties in the near future as temperatures increase in these regions in response to climate changes (Ergon et al. 2018; Helgadóttir et al. 2018). This has created an interest, amongst commercial breeders of perennial ryegrass, in extending the production of this crop further North (Solberg et al. 1994; Helgadóttir et al. 2018). However, there are challenges to this endeavour, including the presence of high levels of GxE interactions, which have previously been documented in perennial ryegrass (Grogan and Gilliland 2011).

These GxE interactions include rank changes, i.e., environment-specific ranking of accessions in respect to productivity (Grogan and Gilliland 2011) and large GxE interactions at year and field level (Fè et al. 2016). Thus, greater insight into the extent of GxE, in perennial ryegrass, in Nordic and Baltic regions may benefit future breeding efforts toward more robust perennial ryegrass germplasm. Few studies have previously investigated GxE in perennial ryegrass and general questions remain about the performance of GxE models in diverse samples. Hence, this study aims to: (i) investigate and assess the performance of perennial ryegrass accessions in a multi-environment field trial, to improve biomass yield and persistence traits, (ii) determine the extent of GxE interactions in perennial ryegrass to assess their potential for adaptation to extreme environments at the margin of the species’ distribution, and (iii) detect adaptive variants which confer significant advantages either across the tested environments, or under specific environmental conditions.

## 2. Material And Methods

### 2.1 Plant material and phenotyping

A total of 264 diploid perennial ryegrass accessions were included in this study. These accessions consisted of advanced cultivars, wild/semi-wild accessions, and landraces obtained from gene banks worldwide, including the Nordic Genetic Resource Center (Rognli et al. 2018). These accessions were multiplied to obtain enough seeds for regular field trials. For the study, a diverse set of accessions was chosen, in respect to geographic origin, which spanned Central Europe, Northern Europe, Russia, US and Japan (Supplementary Table 1). The experiment was conducted across 2014-2016 in Norway (60°45’ N, 11°12’ E), Estonia (58°45’ N, 26°24’ E), Sweden (55°56’ N, 13°6’ E), Lithuania (55°23’ N, 23°52’ E), Iceland (64°9’ N, 21°45’ W) and Denmark (55°20’ N, 12°23’ E). Replicated field trials were established in late-spring or summer in 2014, in Norway (20^th^ of June), Estonia (2^nd^ of August), Sweden (28^th^ of May), Lithuania (10^th^ of June), Iceland (9^th^ of July), and Denmark (25^th^ of June). The elevation, in metres above sea level, or m.a.sl, at the trial sites were as follows: Norway (187 m.a.sl), Estonia (68 m.a.sl), Sweden (65 m.a.sl), Lithuania (33 m.a.sl), Iceland (35 m.a.sl) and Denmark (35 m.a.sl). The following traits were measured in the study: total dry matter yield from three harvests during the growing season (TDM3), spring cover (SpringCover), winter kill (WinterKill). The traits SpringCover and WinterKill are transformed to a scale ranging from 1 to 9, where SpringCover, i.e., percentage of plot covered by the crop, is a measure of the crop’s establishment ability, while WinterKill is measured as the decrease in plot cover from autumn to spring, with 1 for highest survival, 9 for lowest. The data was unbalanced with respect to the number of accessions scored in each country, with the number of accessions varying between 108 and 262 depending on the environment, while across all environments 264 accessions were included. Hence, of the 264 accessions included in the study, a ‘core’ set of 108 accessions were scored in all countries.

### 2.2 Genotyping-by-sequencing

Sequence data was produced using genotyping-by-sequencing (Elshire et al. 2011), with the methylation-sensitive restriction enzyme ApeKI to target the low copy fraction of the genome. Sampling and library preparation followed the protocol described by (Byrne et al. 2013). The accessions were genotyped based on a pooled sample. After alignment of sequencing reads to the reference genome (Nagy et al. 2022) and SNP calling by GATK (McKenna et al. 2010), a total of 380,252 SNPs were obtained. VCF-tools (Danecek et al. 2011) were used to filter the SNP markers that did not meet the following requirements: minDP ≥ 10, min-mean DP ≥ 20, max-meanDP ≥ 100, minQ ≥ 30, minor allele frequency across accessions (MAF) ≥ 0.05, missing rate lower than 20%. After filtering, 151,074 SNPs remained. We then calculated the allele frequency, in each accession *i* where pseudo-counts were used to reduce bias introduced by low read depths: 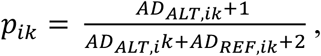, where 𝐴𝐷*_REF,ik_* and 𝐴𝐷*_ALT,ik_* are the depths of the alternate and reference alleles, respectively, in accession *i* at marker *k*. Missing information at a given SNP was imputed as the average allele frequency across accessions.

### 2.3 Genomic relationship matrices

To construct the covariance matrix for additive effects the genomic relationship matrix (GRM) was constructed as 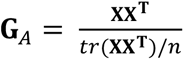 (Vitezica et al. 2017), where **X** is the centered matrix of alternate allele frequency matrix for each accession and marker, *tr* is the sum of the matrix diagonals, and *n* the number of accessions. To construct the covariance matrix for dominance effects, we assumed Hardy-Weinberg equilibrium in the populations, and estimated marker-specific heterozygosity as *H_ik_* = 2*p_ik_ q_ik_*, where 𝑝*_ik_* is the frequency of the alternative allele, and 𝑞*_ik_* is the frequency of the reference allele in accession *i* at marker *k*. Dominance codes were then computed as the residuals of estimated heterozygosity regressed on alternative allele frequency, by extending the approach of (Vitezica et al. 2017). The genomic relationship matrix for the dominance effects was then constructed similarly to the additive GRM.

### 2.4 Environmental data and environmental covariates

Daily weather and climatic variables were extracted for the six trial locations in the period 2014-2016. The information was collected with the R package EnvRtype (Costa-Neto et al. 2021c). Different environmental covariance matrices are constructed for two GxE modelling approaches: (i) reaction norm (RN) and (ii) envirotyping (ET). Environmental similarity was determined based on 18 environmental covariates (Supplementary table 2). For the RN approach we determined the environment means for each environmental covariate x time-window combination.

For the ET approach, daily measurements for each of the environmental covariates were partitioned into quantile intervals (0.01, 0.25, 0.50, 0.99) specific to each time window, intended to characterize environmental zones or envirotypes. The frequency of each envirotype was then determined for each country x year x time-interval combination. For TDM3, the environmental covariates were extracted from the period of the 1^st^ of September, in the year prior to harvest, to the 31^st^ of August the following year (harvest year). For SpringCover and WinterKill, environmental covariates were extracted from the 1^st^ of September in the year prior to when the trait was scored until the 31st of May (year trait is scored). The R-package EnvRtype requires the specification of lower and upper temperature bounds for crop development for the model crop, to which we used experimentally determined temperature bounds as estimated in (Monks et al. 2009). In both approaches the time interval was specified to span a 30-day window. The covariance matrices, denoted by **E**, were computed with the same approach as used for the genomic relationship matrices. For the RN and ET models we defined environments as country-year combinations, hence 12 environments were included in the study (Table 1).

**Table 1:** Number of diploid accessions scored in each country for the traits.

|  | Norway |  | Estonia |  | Sweden |  | Lithuania |  | Iceland |  | Denmark |  |
| --- | --- | --- | --- | --- | --- | --- | --- | --- | --- | --- | --- | --- |
|  | 2015 | 2016 | 2015 | 2016 | 2015 | 2016 | 2015 | 2016 | 2015 | 2016 | 2015 | 2016 |
| **TDM3** | 213 | 213 | 144 | 144 | 108 | 108 | 118 | 118 | - | - | 262 | 262 |
| **SpringCover** | 213 | 213 | 144 | 144 | - | - | 118 | 118 | 142 | 142 | - | - |
| **WinterKill** | - | - | - | 144 | - | 108 | - | 118 | - | - | - | - |

### 2.5 Phenotypic analysis

Linear mixed models were fitted separately in each location to estimate the effects of accession and their interaction with year. Phenotypic records were regressed on accession (fixed), year (random), the accession-by-year interactions (random), spatial effects nested in year (random), and model residuals. The effects of year, the effects of accession-by-year interactions, and model residuals were modelled as independent and identically distributed (*i.i.d.*), following a normal distribution with the same variance for each type of random effect. Random spatial effects differed by countries, due to differences in experimental design: in Denmark, effects of trial and effects of row and column across the field; in Iceland and Lithuania, effects of replicate and effects of row and column in each replicate; in Estonia, Norway and Sweden, effects of trial and effects of row and column in each trial. Effects of trials and replicates were modelled as *i.i.d.*, whereas effects of rows and columns were modelled as normally distributed following a first-order autoregressive correlation structure, by row and column. In each year, accession means were calculated as the sum of the estimated effects of accession and accession-by-year interactions. Due to differences in managements across countries, accession means were centered and scaled in each country x year combination, based on the mean and standard deviation estimated in the core collection of 108 accessions (to account for the unbalanced number of accessions by country).

### 2.6 Genomic models

#### 2.6.1 Main effects of genotype and environment (M1)

Model M1 is a genomic model (GBLUP) which accounts for the main effect of genotype and environment.

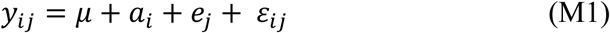

Where 𝑦*_ij_* is the estimated mean phenotypic performance for accession 𝑖 in environment 𝑗 (country-year combination); 𝑒*_j_* the random effect of environment *j* 𝐞 ∼ 𝛮(𝟎, 𝜎*_E_*^2^𝐄); 𝜇 is the intercept, or grand mean; 𝑎*_i_* is the random additive genetic effect of accession *i*, 𝐚 ∼ 𝛮(𝟎, 𝜎*_G_A__*^2^ 𝐆_A_>; while 𝜀*_ijk_* is the error term, 𝜺 ∼ 𝛮(𝟎, 𝜎_𝜺_^2^𝐈), where **I** is the identity matrix.

#### 2.6.2 Additive genetic effect and environment interaction (M2)

The M1 model is extended by adding the GxE interaction term, which is calculated by the Kronecker product of the covariance for the accessions and the covariance for the environments, given as 𝐆***_A_*** ⊗ 𝐄, where 𝑎𝑒 ∼ 𝛮(𝟎, 𝐆**_A_** ⊗ 𝐄 𝜎*_G_A×E__*^2^);

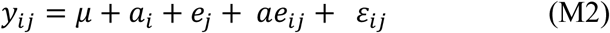

#### 2.6.3 Additive and dominance genetic effects and their interaction with the environment (M3)

In model M3 model we added the random genetic effect of dominance *d*_i_ and its interaction with the environment, where 𝐝 ∼ 𝚴(0, 𝐆*_D_*𝜎*_G_D__*^2^) and 𝑑𝑒 ∼𝚴(0, 𝐺_D_ ⊗ 𝐄𝜎*_G_D×E__*^2^).

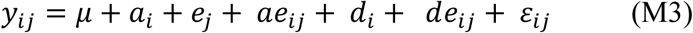

#### 2.6.4 Heritability

Narrow-sense heritability (ℎ^2^) measures the proportion of the phenotypic variance which can be explained by the additive genetic variance: Narrow-sense heritability was calculated based on the variance components from the M3 model as:

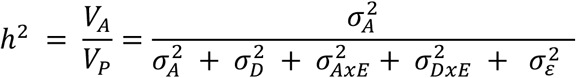

### 2.7 Model comparison and prediction accuracy

Likelihood ratio tests based on restricted maximum likelihoods (REML) were used for model comparison. The likelihood ratios were calculated as: 𝐿𝑅 = −2 [𝑙_0_(𝜃) − 𝑙_1_(𝜃)], where 𝑙_1_(𝜃) is the log-likelihood of the tested model, and 𝑙_0_(𝜃) the log-likelihood of the null model. The likelihood ratio test statistic follows a chi-square distribution, where the degrees of freedom is the difference in the number of parameters (random effects) between the models. The models were tested under three different validation schemes (i) Leave-one-country-out, where the models’ ability to extrapolate to unobserved environments was tested. (ii) Leave-one-random-cluster-out, where accessions are randomly assigned to ten clusters. The validation scheme is used to evaluate model ability to predict unobserved accessions when related genetic material is included in the training-set. (iii) Leave-one-genetic-cluster-out, where we divided accessions into 10 genetic clusters based on genetic similarity. This validation scheme is used to evaluate model performance when the genetic similarity between accessions in the training and test set is low. Genetic clusters were inferred by hierarchical clustering based on Ward’s criterion on the matrix of alternate allele frequencies **X**, with Euclidean distance as distance metric. Prediction accuracy was calculated for each fold as the Pearson correlation between observed accession means and predicted accession means *ŷ_ij_*: 𝐶𝑜𝑟(𝑦_ij_, *ŷ_ij_*). Finally, model performance, i.e. prediction ability, was calculated as the mean correlation across all folds within the given validation scheme.

### 2.8 Genome-Wide-Association study (GWAS)

A linear mixed model approach was applied to identify associations between accession means and genetic variants. Two different GWAS models were fitted: (i) a ‘marginal’ GWAS model to detect QTL with consistent effects in the suite of environments included in the study; and (ii) a ‘conditional’ GWAS to discover environment-specific QTL effects. To account for population structure, we included the fixed effects of the first three principal components from a principal component analysis (PCA) of the centered allele frequency matrix **X**. In marginal GWAS, the fixed effect of the tested SNP was 𝑠_i_𝛼, where 𝑠_i_ is the alternate allele frequency at the tested SNP for accession *i* and *α* is the effect of the alternate allele on the phenotype:

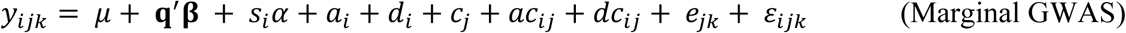

where **q** is the vector of the principal components for accession *i* and **β** is the vector of associated effects; 𝑐_j_ is the fixed effect of country *j*; 𝑎_i_ and 𝑑_j_ are the additive and dominance genetic effects as described above; 𝑎𝑐_ij_ and 𝑑𝑐_ij_ are interactions between additive genetic effects and country, 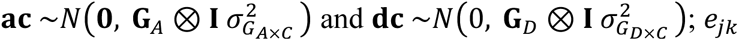 is the random effect of environment *jk* (combination of country *j* and year *k*); 𝜀*_ijk_* is the model residual.

To detect genetic variants in linkage with QTL which have environment-specific effects, we extended the marginal GWAS model by including the interaction between SNP and country:

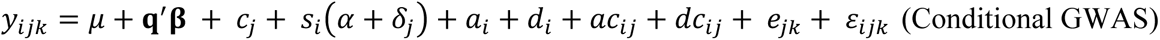

where 𝛿_j_ is effect of the SNP in country 𝑗. Significance was assessed by a Wald test. To correct for multiple testing, statistical significance was assessed by the false discovery rate (FDR) (Benjamini and Hochberg 1995).

### 2.9 Software

All models were constructed in R 4.2.0 (Development Core Team) within high-performance computing cluster GenomeDK. The R-package MM4LMM version 3.0.2 (Laporte et al. 2022) was used to extract model likelihoods, model validation and to run the GWAS analysis. The R-package EnvRtype version 1.1.1 (Costa-Neto et al. 2021c) was used to extract environmental covariates and construct the environmental covariance matrices for the two modelling approaches. For the phenotypic analysis we used the software ASREML-R version 4.2 (Butler et al.). The R-package Dendextend was used to infer genetic clusters (Galili 2015).

## 3. Results

### 3.1 Perennial ryegrass accessions are highly diverse and exhibit minimal population structure

Generally, the germplasm included in this study was quite diverse, with the first PC accounting for less than 2 % of genomic variation. Additionally, we observed rapid LD-decay, with the genome-wide average LD decreasing to below r^2^ < 0.2 after 200 bp (Figure 1). This is indicative of a germplasm being highly genetically dissimilar, and that alleles at loci in close physical proximity are largely randomly associated. There was a moderate correlation (0.62) between the additive and dominance genomic relationship matrices for the off-diagonal elements, which indicates that there is some collinearity between the additive and the dominance effects. We identified 10 genetic clusters based on hierarchical clustering, which were used for the leave-one-genetic-cluster-out validation scheme (Figure 1).

**Figure 1:**
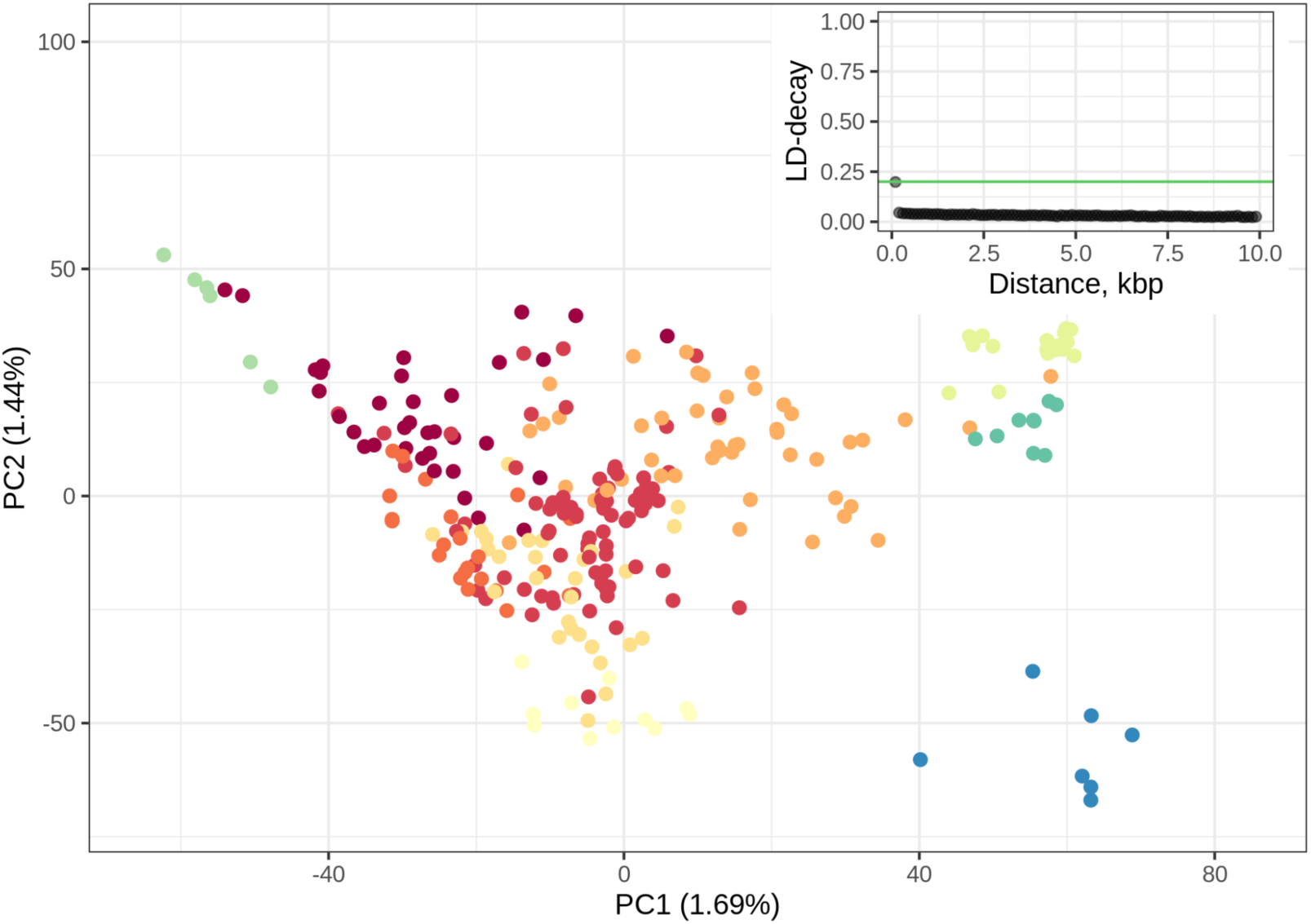
PCA of the 264 perennial ryegrass accessions based on their additive allele dosage. Colours indicate which genetic cluster the accessions have been assigned, based on hierarchical clustering. Linkage disequilibrium (LD) decay is shown in the upper-right corner, as a function of distance in kilobase pairs (kbp). Threshold shown at LD, *r*^2^ = 0.20.

### 3.2 Observed differences among accessions depend on additive and dominance genetic effects and their interactions with climate

Estimated variance components indicate that the additive genetic components contributed most to the phenotypic variance, regardless of trait or modelling approach, RN or ET (Figure 2). The additive genetic component amounted to 37-41 % of the variance for TDM3, 41-43 % for SpringCover and 28-33 % for WinterKill (Figure 2). For TDM3, the interaction between the dominance genetic effect and environment (DxEnv) constituted about 17% of phenotypic variance, while the interaction between the additive genetic effect and environment (AxEnv) constituted about 15-18%. Therefore, the two forms of GxE had approximately equal contributions to the phenotypic variance in TDM3. This indicates that heterozygosity, and in turn genetic variation within accession, may be important for environment-specific interactions and local adaptation. For SpringCover and WinterKill environment-specific interactions are better captured by the interaction between the additive genetic component and the environment (AxEnv), with this component accounting for 25-31% of the variance in SpringCover and 22-26% of the variance in WinterKill, whereas the interaction between the dominance component and the environment (DxEnv) contributed with 0.4-0.9% of the variance in SpringCover and 11-14% in WinterKill. The dominance genetic component’s contribution to phenotypic variance was negligible in SpringCover and WinterKill, while in TDM3 it accounted for approx. 14% of the phenotypic variance. The environmental component contributed minimally to the phenotypic variance for all traits, as accession means were centered and scaled in each environment (based on means and standard deviations in the core collection of 108 accessions), to avoid confounding by management effects. Lastly, heritability estimates did not differ markedly between the RN or ET modelling approach for any of the traits, with all traits showing moderate narrow-sense heritabilities (Table 2).

**Figure 2:**
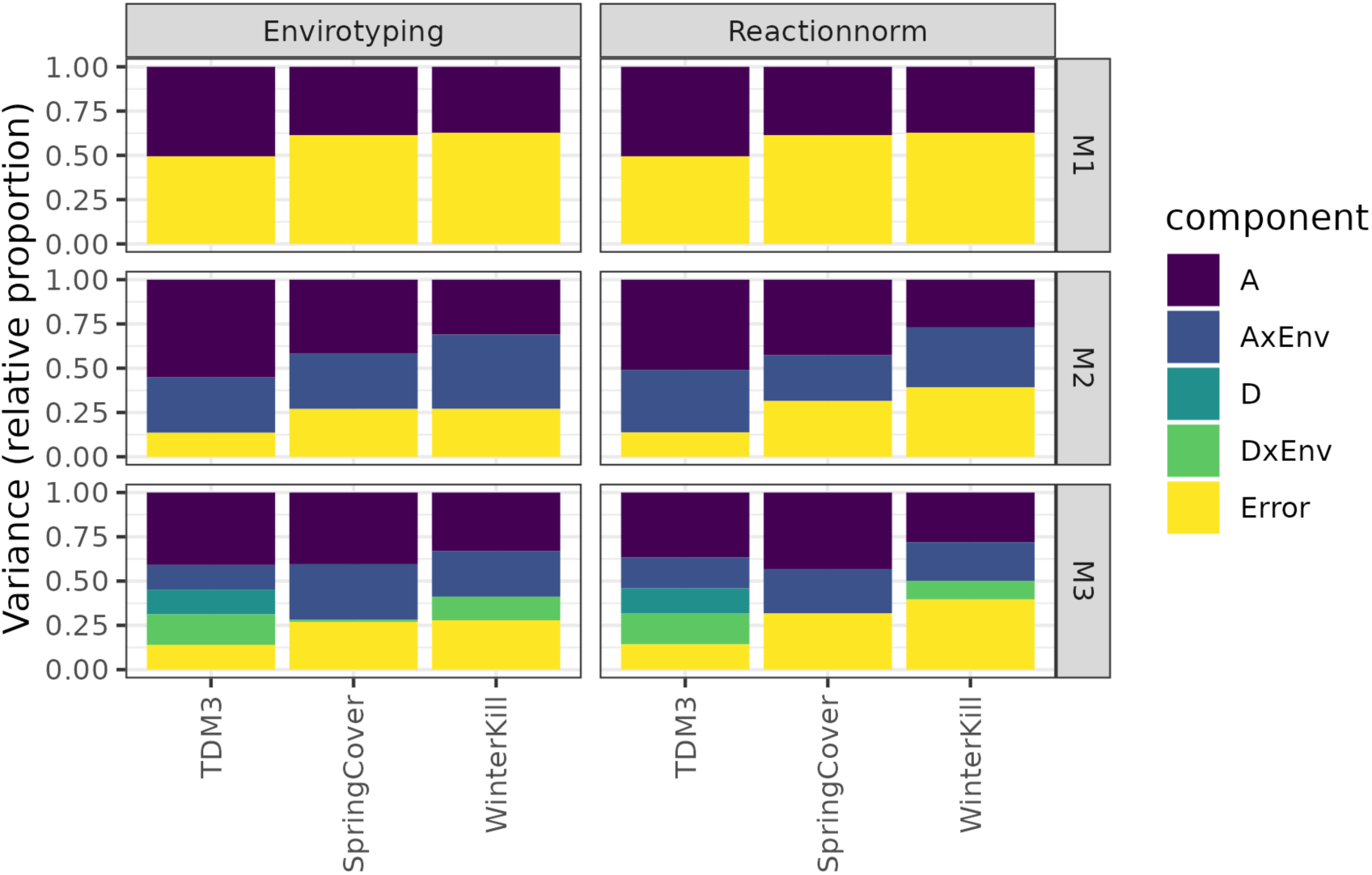
Relative proportion of variance explained by the different model components, excluding the environmental component. M1: main effects of genotype and environment; M2: main effects of genotype (additive) and their interactions with environment; M3: main effects of additive and dominance genetic effects and their interactions with the environment. A: additive genetic variance component, AxEnv: variance component for the interaction between additive genetic effect and environment, D: dominance genetic variance component, DxEnv: variance component for the interaction between the dominance genetic effect and the environment, where error is the variance component for the residuals.

**Table 2:** Narrow-sense heritability (h^2^) for all traits, as estimated by model M3 (A + Env + D + AxEnv + DxEnv) for the Reaction-norm (RN) and Envirotyping (ET) approaches.

|  | **TDM3** | **SpringCover** | **WinterKill** |
| --- | --- | --- | --- |
| **RN** | 0.37 | 0.43 | 0.28 |
| **ET** | 0.41 | 0.41 | 0.32 |

We compared the fits of the nested models M1, M2 and M3 (Table 3). Results show that the addition of an interaction term between the additive genetic component and environment significantly improves model fit for all traits. The addition of a dominance term and an interaction term between the dominance effect and the environment, led to significant model improvement for TDM3, but not for SpringCover and WinterKill. These results suggest the presence of QTL with environment-specific effects which may confer adaptation to specific climatic conditions. Additionally, results indicate that environment-specific marker effects on TDM3 may be mediated by dominance or genetic diversity, in addition to additive genetic effects. In contrast, for SpringCover and WinterKill, results indicate that environment-specific marker effects are mainly mediated by additive genetic effects, and the interaction between additive genetic effects and environment. Independent of the trait, we find that the RN and ET models do not differ in their ability to capture GxE interactions.

**Table 3:**
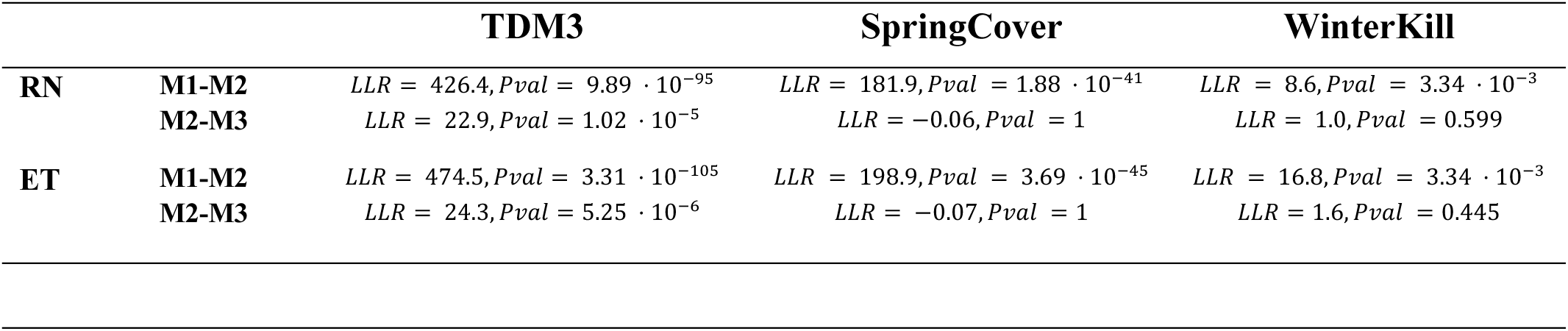
Log-likelihood ratio (LLR) statistics and P-values (Pval) for model comparisons, from likelihood-ratio tests. M1: main effects of genotype and environment; M2: main effects of genotype and their interactions with environment; M3: main effects of additive and dominance genetic effects and their interactions with the environment.

### 3.3. Accounting for interactions between additive effects and climate variables improves genomic prediction accuracy

Variance component estimates and model comparison with likelihood ratio tests indicate that including GxE interactions might improve model performance, i.e., prediction ability (PA). To determine if GxE interactions improves evaluation of accessions in diverse samples and environments, the prediction accuracy of our models is evaluated under three validation schemes: i) unobserved environments (leave-one-country-out validation), ii) unobserved germplasm, grouped based on relatedness (leave-one-genetic-cluster-out validation), and iii) unobserved germplasm, randomly grouped (leave-one-random-cluster-out validation). PA in unobserved environments (leave-one-country-out validation scheme), increased with the inclusion of GxE interactions for TDM3, but not for SpringCover and WinterKill. PA for TDM3 increased by up to 4% when accounting for additive GxE, whereas for SpringCover, PA decreased by 2-4% depending on the modelling approach, while for WinterKill inclusion of additional model terms, with M3, led to an 8% decrease in PA with the ET approach and a 24% decrease with RN (Table 4). The lower PA when GxE terms was included for SpringCover and WinterKill, may suggest the presence of QTL with highly environment-specific effects which would limit extrapolation of marker effects to novel environments. Results of leave-one-genetic-cluster-out validation show that for TDM3, model M1 cannot accurately extrapolate to novel germplasm, with PA effectively around zero (Table 4). However, addition of an interaction term between the additive genetic effect and environment (AxEnv) improves PA for yield by up to 11%. Further extension of the model by inclusion of additional terms (D and DxEnv) further increases PA by up to 6%. For SpringCover and WinterKill, we observe similar improvements in PA, which increased by up to 21-22% and 7-9%, respectively, when accounting for AxEnv. These results support the presence of QTL with environment-specific effects which covary with the environmental covariates used to characterize the environments. These moderate improvements in PA further indicate that despite the accessions being genetically diverse, the models are capable of extrapolating GxE interactions to genetically distinct clusters of accessions. For the prediction of unobserved germplasm, randomly assigned into clusters (leave-one-random-cluster-out validation scheme), the PA of the M1 model is much improved for all traits compared to the PAs observed in leave-one-genetic-cluster-out validation. However, limited improvement in PA is observed when extending the M1 with GxE terms under this validation scheme compared to the leave-one-genetic-cluster-out scheme, with PA increasing by up to 2% (TDM3), 5-6% (SpringCover) and 2-3% (WinterKill). The large improvement in PA for the leave-one-genetic-cluster-out validation compared to the leave-one-random-cluster-out validation may suggest that the additive genetic effects alone may be less useful when predicting in novel and genetic dissimilar germplasm, and environment-specific marker effects may provide useful information in these challenging situations. Our results show that inclusion of GxE may lead to improved PA in one validation scheme but confer no improvement or potentially reduce PA in another (Figure 3), e.g., for SpringCover and WinterKill where we observed improved PA in novel germplasm but lower PA in novel environments. Neither of the two GxE modelling approaches (RN and ET) consistently outperformed the other under the three validation schemes. For lack of clear evidence of either modelling approach being superior, we will for downstream analyses use the widely used RN modelling approach.

**Figure 3:**
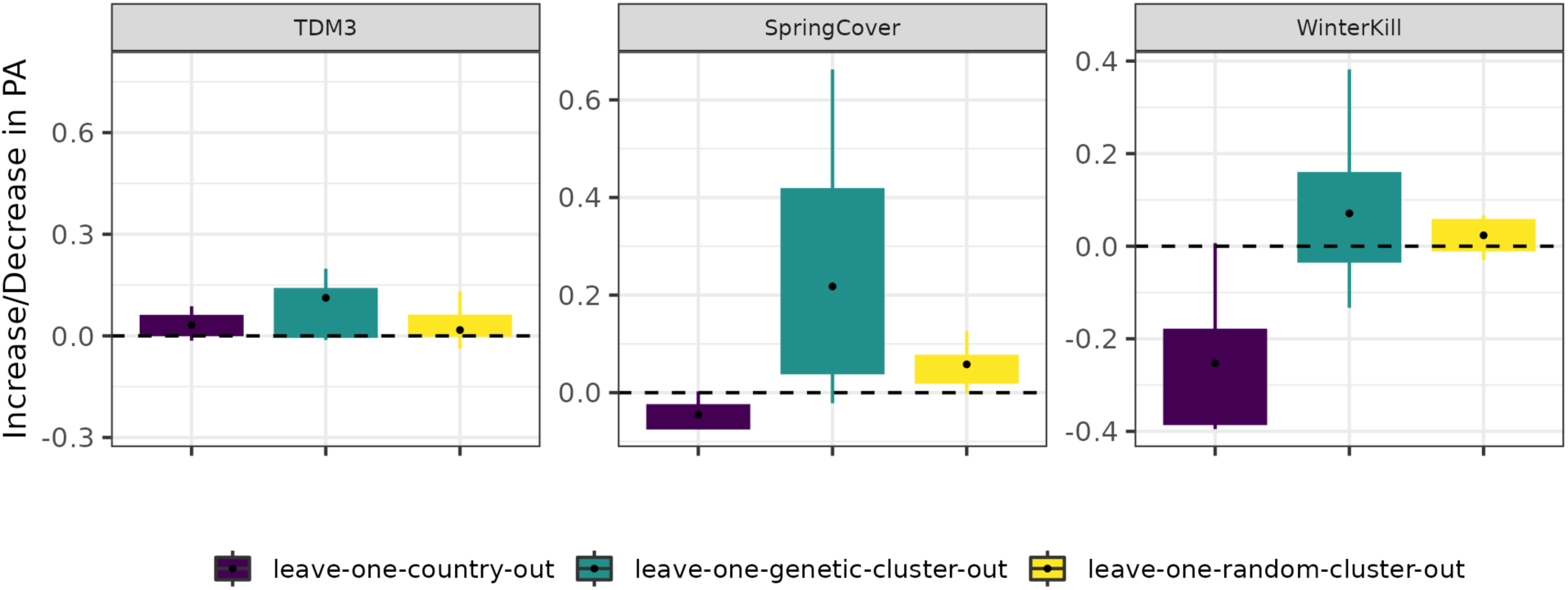
Increase/decrease in PA from M1 to M3 shown for each validation scheme, specifically for the reaction norm approach. The average PA for each validation scheme is denoted by a black dot.

**Table 4:**
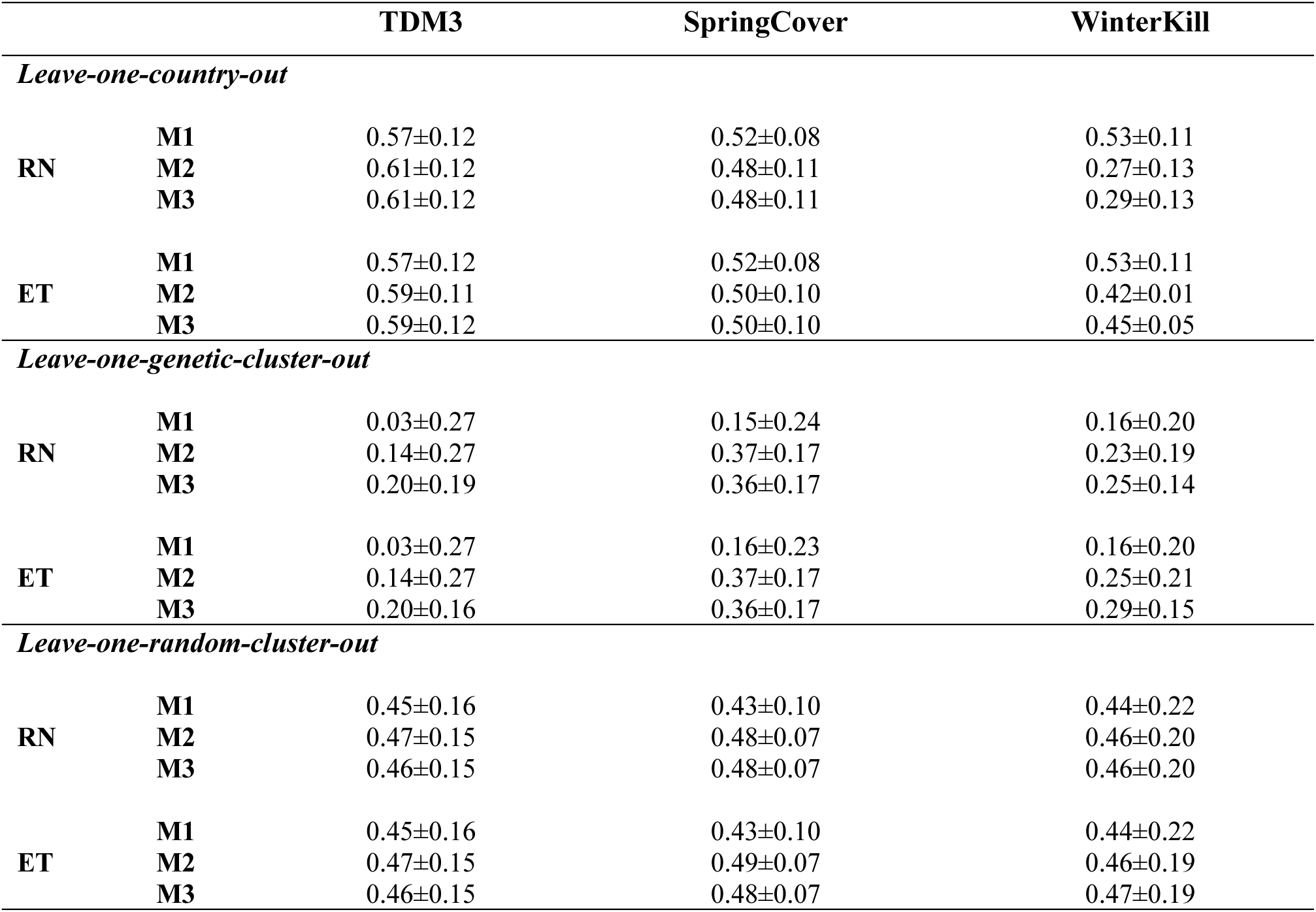
Mean prediction accuracy and standard deviation across validation sets, in the three validation schemes (Leave-one-country-out, Leave-one-genetic-cluster-out, and Leave-one-random-cluster-out) for the two modelling approaches: Reaction-norm (RN) and Envirotyping (ET). M1: main effects of genotype and environment; M2: main effects of genotype and their interactions with environment; M3: main effects of additive and dominance genetic effects and their interactions with the environment.

### 3.4 Genomic prediction accuracy depends on countries

Environments included in this study was diverse and each uniquely challenging for the crop, which can increase the difficulty of capturing any GxE interaction. We compare PA for each country to determine whether certain environments constitute a larger challenge for our models (Figure 4). We note that PA for TDM3 and SpringCover is relatively stable across environments, while PA for WinterKill varies greatly across countries. These results indicate that WinterKill may be a trait which is particularly challenging to predict in novel environments. However, it is uncertain whether this is due to sparse sampling of environments (i.e., too few across a large area), highly environment-specific effect marker effects, and/or selected environmental variables not correlating well with phenotypic performance in specific environments.

**Figure 4:**
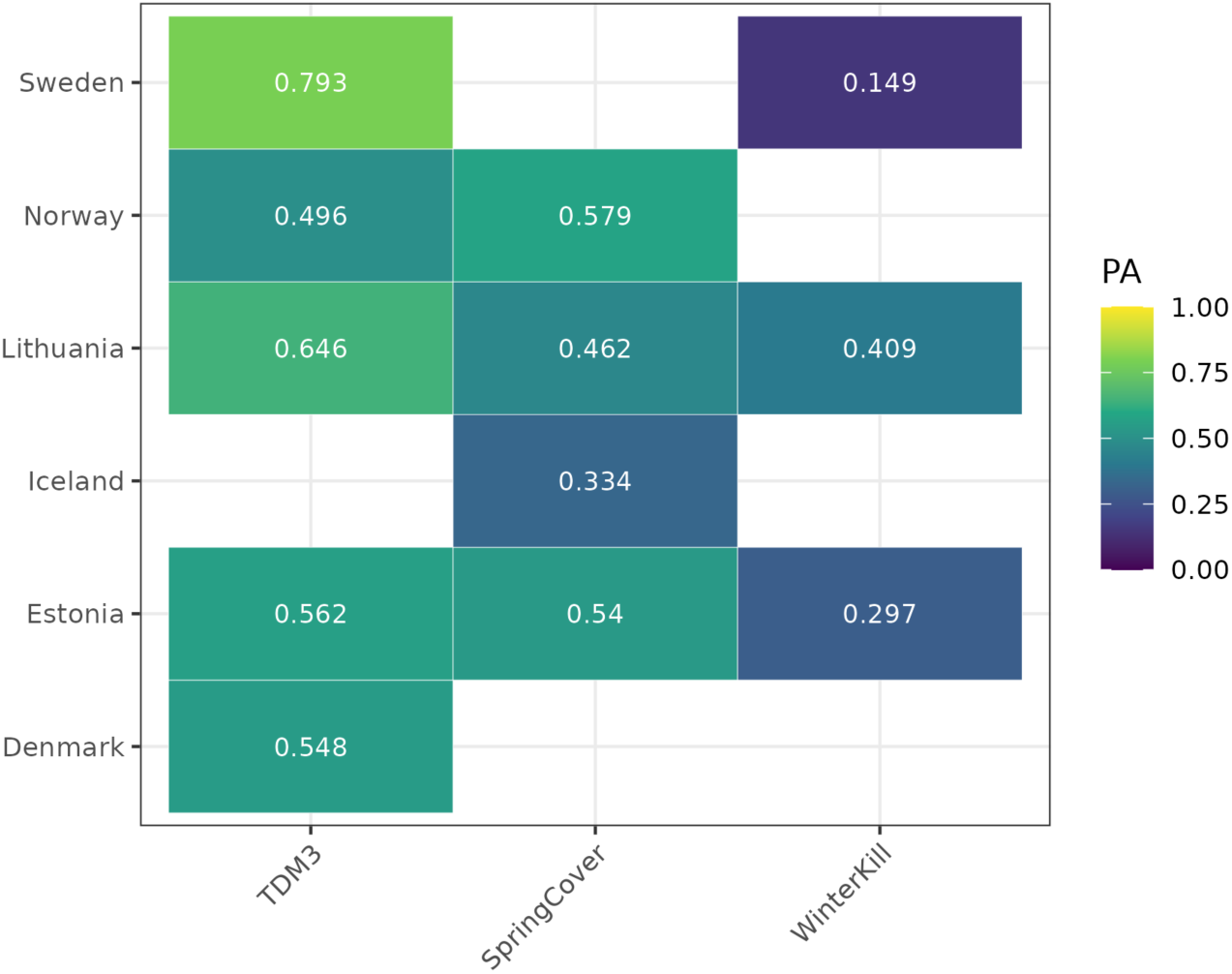
PA shown for each country in the leave-one-country-out validation scheme, where the GxE effects are modelled with the reaction norm (RN) approach. PA is shown for the M3 model which includes the additive and dominance genetic effects and their interaction with the environment.

### 3.5 Genomic prediction models predict different components of phenotypic variability with varying accuracy

Validations show improved PA when accounting for GxE for some traits and under certain validation schemes, while for others there is a cost to including GxE (Figure 3). Here we investigate which model components constitute a greater challenge for our GP models. We compare the correlation between accession means 𝑦_ij_ (for any accession *i* in environment *j*), and the estimated effects for each model component i.e., additive genetic effects (*â_i_*), dominance genetic effects (*d̂_i_*), interactions between additive genetic effects and environment (*âe_ij_*), and dominance genetic effects and environment (*d̂e_ij_*). Correlations between component effects and accession means are calculated in validation sets. This analysis highlights the components which are more challenging for our models to extrapolate and/or interpolate (Figure 5). The genetic effects (*â_i_* and *d̂_i_*) are generally well predicted for the leave-one-country-out validation scheme, with 𝑐𝑜𝑟= (*y_ij_*, *â_ij_*) (averaged over validation sets) ranging from 0.49 to 0.59, while 𝑐𝑜𝑟=(*y_ij_*, *d̂_ij_*) ranged from 0.39 to 0.56. When interpolating to novel germplasm, where random accessions are assigned to the validation set, the genetic effects are still reasonably well predicted, with 𝑐𝑜𝑟= (*y_ij_*, *â_ij_*) ranging from 0.41 to 0.43, while 𝑐𝑜𝑟= (*y_ij_*, *d̂_ij_*) ranged from 0.19 to 0.37. However, in the leave-one-genetic-cluster-out validation scheme, extrapolation of genetic effects fails, with 𝑐𝑜𝑟= (*y_ij_*, *â_ij_*) ranging from 0.07 to 0.16 while 𝑐𝑜𝑟= (*y_ij_*, *d̂_ij_*) ranged from -0.05 to 0.14. The leave-one-genetic-cluster-out validation scheme is furthermore characterized by large standard deviations indicating that for certain genetic clusters the genetic effects may be well predicted, whereas for other clusters 𝑐𝑜𝑟= (*y_ij_*, *â_ij_*) and 𝑐𝑜𝑟= (*y_ij_*, *d̂_ij_*) are either effectively zero or negatively correlated with phenotypic performance (Figure 5). Correlations between accession means and estimated GxE interactions show that coefficients *âe_ij_* and *d̂e_ij_* are generally more challenging to predict under the leave-one-country-out validation scheme, as suggested by Figure 4, which show a consistently larger cost to accounting for GxE terms in novel environments, as shown by a decrease in PA from the M1 to M3 model. Together these results underlines that the lower PA for the leave-one-genetic-cluster-out validation scheme, specifically with M1 model, compared to the other validation schemes is due to the models not being able to extrapolate genetic effects (*â_i_* and *d̂_i_*) to novel, and genetically distinct, germplasm. Our results further indicate that under the circumstances where the main genetic effects are not well predicted then there is a greater benefit to accounting for GxE, as shown by larger increases in PA under the leave-one-genetic-cluster-out scheme compared to the leave-one-random-cluster-out where main genetic effects are more useful to predict accession means.

**Figure 5:**
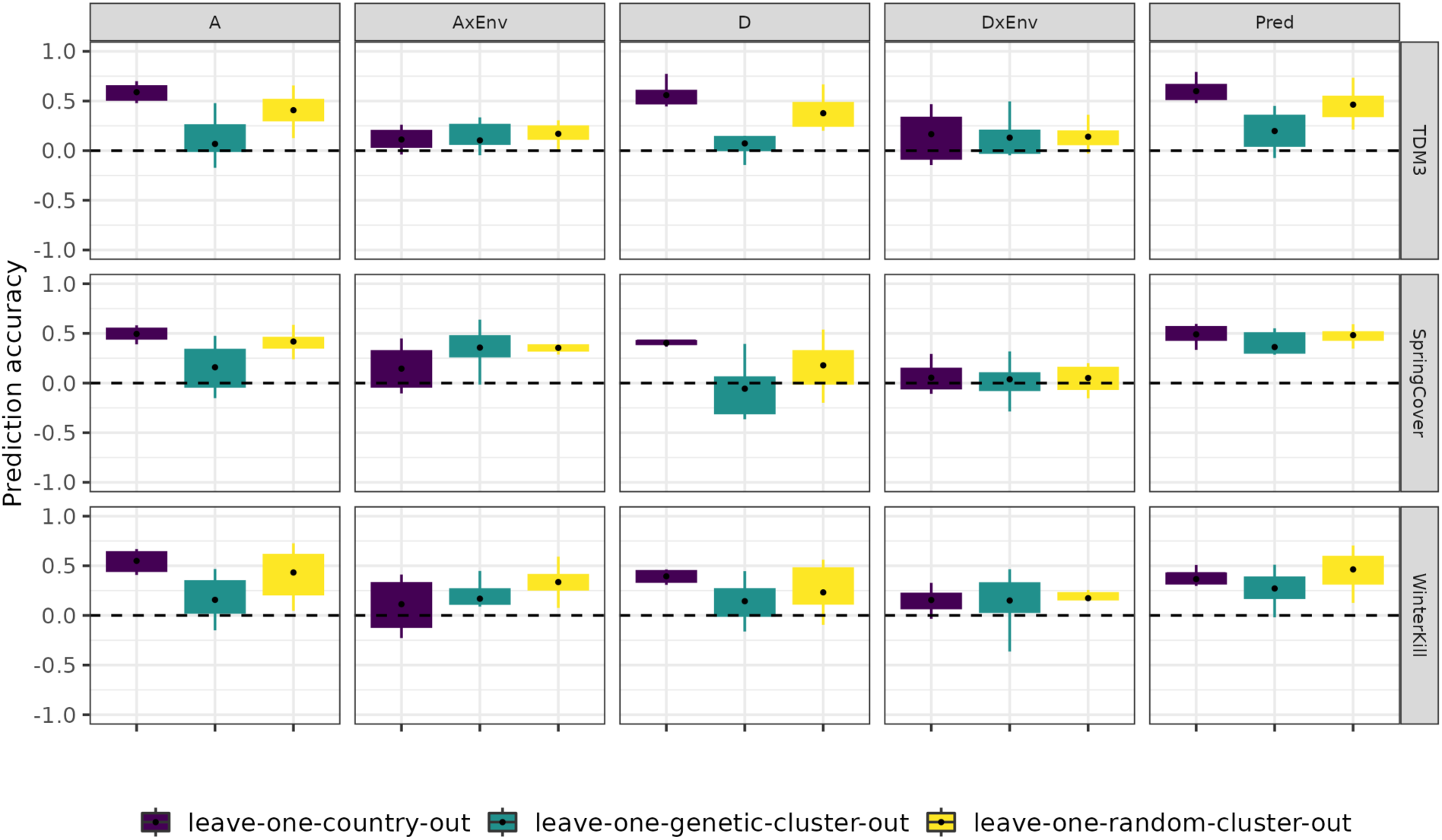
Correlations in left-out folds for the three validation schemes (Leave-one-country-out scheme, Leave-one-genetic-cluster-out and Leave-one-random-cluster-out) from model M3 fitted with the reaction-norm approach. The following correlations are shown: Pred., Observed accession means vs. predicted accession means; A, Observed accession means vs. Additive breeding values coefficients; D, Observed accession means vs. Dominance breeding values coefficients; Env, Observed accession means vs. Environment coefficients; AxEnv, Observed accession means vs. AxEnv coefficients; DxEnv, Observed accession means vs. DxEnv coefficients.

### 3.6 Genome-wide association studies detected marginal and country-dependent marker effects

We conducted GWASs to detect genetic variants with marginal effects, i.e. variants with environment-independent effects, and conditional effects, i.e. environment-dependent effects (Figure 6).

**Figure 6:**
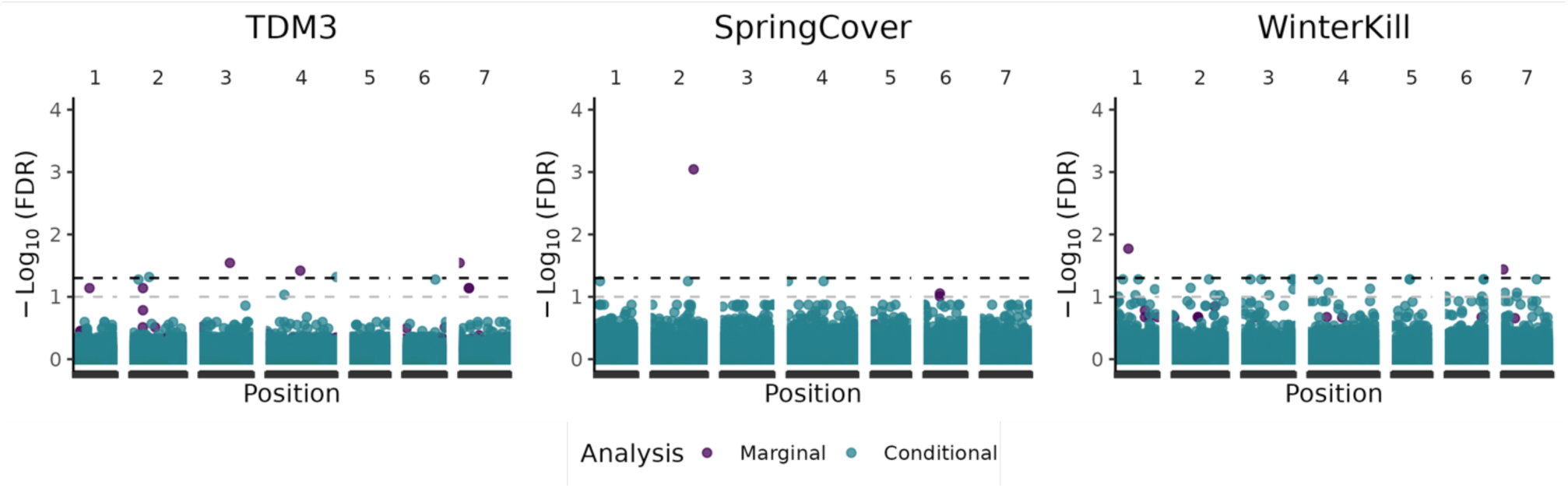
GWAS results for a marginal and conditional GWAS analysis. Black line denotes the significance threshold (0.05) adjusted with FDR (False-discovery rate), while grey line denotes the significance threshold (0.10) adjusted with FDR.

Previous analyses indicated extensive additive genetic effects (Figure 2 and Table 3) and moderate narrow-sense heritabilities (Table 2) for all traits, suggesting that a large proportion of the phenotypic variation may be described by additive genetic variation. However, few significant QTL were detected by the marginal GWAS. These results suggest that the investigated traits are polygenic, and are characterized by many small-effect QTL. Genome-wide analyses of GxE, i.e., likelihood-ratio tests (Table 3) and GP validation (Table 4), indicated the presence of environment-specific marker effects. Few conditional effects were highly significant (FDR ≤ 0.05) for TDM3. For SpringCover, there was some evidence for potential QTL (FDR ≤ 0.10) conferring a disadvantage in specific environments (Figure 7). Similarly, for WinterKill we detected potential QTL which may confer an advantage in Lithuania but not in Sweden (Figure 7).

**Figure 7:**
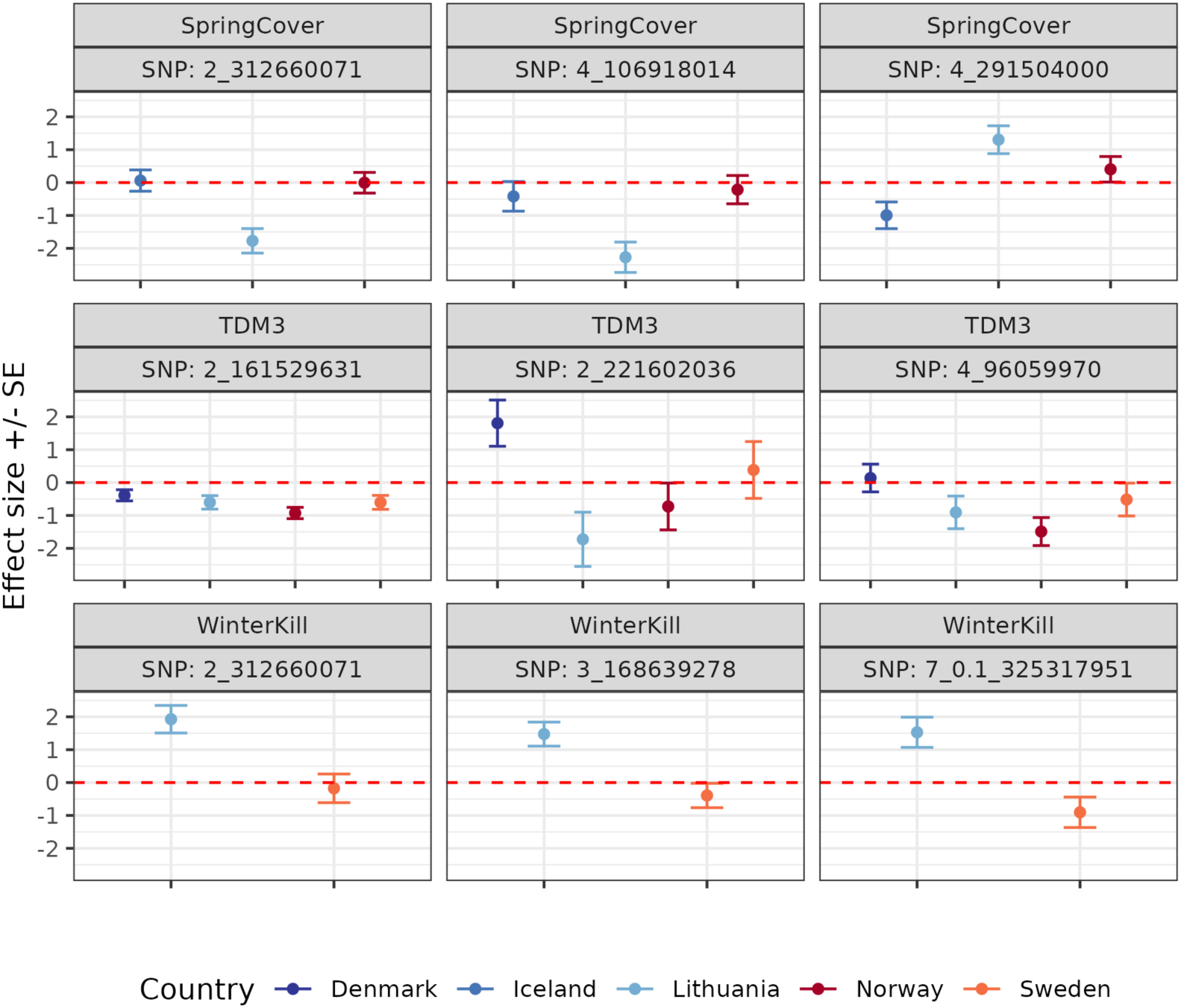
Effect estimates ± standard errors (SE) for the SNPs with the most significant P-values from the conditional GWAS.

For both the marginal and conditional GWASs we found few significant QTL, which may be due in part to the germplasm being genetically dissimilar as demonstrated by limited LD (Figure 1). Hence, only common and large-effect QTL would be detected by our models. Alternatively, it is possible that the underlying genetic architecture is highly polygenic, both in regard to the additive genetic effects and the additive GxE effects, implying that most QTL have small effects and thus challenging for the models to detect.

## 4. Discussion

### 4.1 Diverse accessions contain unique genetic variation important for broadening the genetic base of perennial ryegrass

This study included a geographically and genetically diverse germplasm, to enhance the probability of including and identifying genetic variation that may confer an advantage under the environmental conditions experienced in the Nordic and Baltic countries. Utilising genetically diverse accessions is favourable in prebreeding, as less advanced breeding material, or potentially unselected material, may be enriched for higher genetic diversity and may contain novel adaptive variants not present in current breeding programs, which may be introgressed into more advanced breeding material for enhanced phenotypic performance under biotic and abiotic stressors as demonstrated in previous studies (Meseka et al. 2013; Singh et al. 2021). The present study sought to quantify and estimate the extent of GxE interactions in environments that are at the limit of the species’ distribution. GxE interactions are environment specific, hence while previous studies have investigated GxE interactions in perennial ryegrass, most of these studies have investigated GxE in temperate regions, which are currently more suitable for its production e.g., in central and southern Europe (Fois et al. 2021), Denmark (Fè et al. 2015, 2016) and Ireland (Grogan and Gilliland 2011). Consequently, the findings of these studies are presumably not indicative of the level of GxE interactions that are present, or occur, in the morer extreme environments experienced in the northern regions, investigated in the present study. GxE interactions in environments comparable to those investigated in this study have previously been documented in perennial ryegrass, by (Helgadóttir et al. 2018) who found that commercial cultivars are not widely adapted to the environmental conditions across the Nordic and Baltic regions, which underlines the importance of including unselected material in future breeding efforts. Additionally, the accessions included in the study have been collected from vastly different geographic origins, which is a benefit as local adaptation to some of these environments may confer an advantage for the environments tested in the current study. Hence, if we instead had included genetically similar accessions, we would have had to constrain us to only include highly related individuals (from similar geographic backgrounds), which would have been a limitation.

### 4.2 Genotype x environment interactions pose challenges in diverse samples of environments and accessions

In the current study we compared a standard reaction-norm approach, as demonstrated by (Jarquín et al. 2014) to a more recent envirotyping approach as demonstrated by (Xu 2016) and later extended by (Costa-Neto et al. 2021b, a). In accordance with the findings of previous studies we found phenotypic variation to be highly affected by GxE interactions for both ordinal traits, i.e., SpringCover and WinterKill, as well as yield traits. The inclusion of interactions between additive genetic effects and environments (AxEnv) led to moderate improvements in model fit and prediction accuracy for all traits. Inclusion of dominance and dominance GxE terms further improved prediction and model fit. Previous studies, in hybrid maize, have similarly documented the importance of dominance genetic effects in regard to yield production, with (Rogers et al. 2021) demonstrating that the dominance genetic effect was a more effective predictor of yield performance than the additive genetic effect. However, when model performance was evaluated based on their ability to predict the phenotypic performance of novel germplasm, where training and validation sets were genetically more similar (leave-one-random-cluster-out) or when predicting the performance of observed germplasm in novel environments (leave-one-country-out), there were more limited improvements in prediction accuracy with the inclusion of GxE terms. Previous GxE studies (e.g., in wheat) experienced more marked improvements in prediction accuracy when accounting for GxE, with some studies citing increases in prediction accuracy by up to 82% when predicting yield performance of unobserved lines, modelled with a reaction-norm approach (Jarquín et al. 2017), while (Heslot et al. 2014)reported an increase in mean prediction accuracy of approximately 25-28% for yield production in winter wheat. It should be noted that the study by (Heslot et al. 2014)utilized crop growth models that incorporate developmental stage specific environmental covariates (stress covariates), and hence the environmental covariates included in that study was presumably more physiologically relevant than the environmental covariates included in this study. Generally, GxE studies cover less heterogeneous areas than those investigated in this study, e.g., Northern France (Jarquín et al. 2014) and France (Heslot et al. 2014). Additionally, these studies also investigated GxE interactions for across a denser spatial and temporal grid, with the study by (Heslot et al. 2014) including 44 environments (location-year combinations) from 2006 to 2011, while the study by (Jarquín et al. 2014) investigated 134 locations across eight years. The importance of employing a dense sampling approach is further underlined by (Rogers and Holland 2022), where they noted that more sparsely sampled areas suffered higher reductions in prediction accuracy as these environments were less represented by the environments included in the training set. This is potentially also why we observed reduced PA for yield in Iceland, which is environmentally most dissimilar to the other environments (Figure 4). Therefore, more effective extrapolation of GxE might be achieved with denser environmental grids, even with large differences among environments. Additionally, in the current study, we did not perform feature selection of the environmental covariates. However, depending on the trait and environmental conditions at the trial sites it is possible that certain environmental covariates will not covary with the tested traits, thus inclusion of these environmental covariates will introduce noise. As such, this analysis may potentially benefit from feature selection of the environmental covariates which has been shown to lead to improved PA in other studies (Montesinos-López et al. 2023). However, the dissimilarity of the environments tested in this study, may challenge feature selection as the relevance of different environmental covariates may vary across trial sites. Further suggesting that a less sparse spatial grid is required for further model refinement with e.g. feature selection.

### 4.3 Genetic diversity and rapid LD decay in perennial ryegrass limit the accuracy of genomic prediction

The improvements in PA, observed for certain validation schemes, are more modest than those demonstrated in previously mentioned studies. This is to be expected as the accessions are genetically diverse, and include unselected material where genomic relationships between accessions are limited. This is further compounded, as we inferred genetic clusters based on genetic similarity, resulting in increased genetic dissimilarity between the accessions in the training and validation sets. Consequently, the lack of population structure and low levels of genetic similarity between accessions in the training set and the validation set might have limited our ability to efficiently extrapolate both additive genetic effects and GxE interactions to unobserved accessions. This is supported by the findings of (Pembleton et al. 2018), which demonstrated that prediction accuracies were negatively affected by the inclusion of genetically distinct subgroups of perennial ryegrass, and that prediction accuracies could be enhanced by removing genetic material that is genetically distinct from the majority of the germplasm contained in the training sets. Consequently, while low genetic similarity, in general, is a challenge across species, it is possible that there is a particularly large cost to including genetically dissimilar material when performing genomic prediction in perennial ryegrass, a species characterized by rapid LD decay of less than 3 kb (Ponting et al. 2007). In our study we found rapid declines in linkage disequilibrium (LD), with LD decay within 200 bp (Figure 1), hence we underline the importance of including genetically similar germplasm for genomic selection in perennial ryegrass. However, focusing on advanced and highly related individuals would have limited the probability of identifying novel adaptive genetic variation which may be introgressed into advanced breeding material and in turn may enrich the genetic diversity in perennial ryegrass breeding populations.

### 4.4 Applications for Breeders: perennial ryegrass and its future in the Northern regions

For breeders to be able to produce perennial ryegrass in more northern regions it is of importance to identify genetic material that can survive the cold and variable climates experienced in these regions. In the present study, we were able to identify genetic variants in linkage with QTL that might be associated with increased winter survival. These variants appear to have environment-independent effects across the environments where WinterKill was scored. The presence of extensive GxE (Figure 2 and Table 3) suggests that it may be advantageous to breed for local adaptation to specific environmental conditions or countries, as introgression of QTL with environment-specific effects may be advantageous in certain conditions while deleterious in others. Furthermore, depending on the stochasticity of the environmental stressors, at the intended production area, considerations regarding whether to select for increased GxE interactions should be made. As environments with reliable environmental cues, e.g. seasonal changes in day-length or temperature reliably signalling future selective environments, there is an increased chance of GxE interactions improving population persistence under stressing conditions, whereas if the environment is highly stochastic and environmental cues for acclimatisation are unreliable such interactions may be deleterious for survival (Reed et al. 2010). Hence, if the environments at the intended production area are characterised by variable and increasingly unpredictable weather patterns it may be more beneficial to select for accessions with smaller GxE interactions.

### 4.5 Conclusions

Our study highlighted extensive GxE in perennial ryegrass for the environments experienced in the Nordic and Baltic regions. We found no clear evidence for different modelling approaches surpassing the other (either reaction norm based on linear response to environmental variables, or envirotyping, based on non-linear responses). The environments included in this study were highly diverse, and we observed that the statistical models were challenged in modelling the environmental effect on phenotypic performance for the majority of the traits, presumably because of sparsity in the spatial grid. While there was some benefit in accounting for GxE when predicting the performance of unobserved and genetically dissimilar germplasm, these moderate improvements in prediction were not observed for validation schemes where the additive genetic effects were most useful to predict phenotypic performance (leave-one-random-cluster out validation). Despite rapid LD decay in our diverse panel of accessions, we were able to detect several candidate QTL associated with multiple traits, including QTL with country-specific effects. Together, our results suggest that there is adaptive genetic variation in the perennial ryegrass diversity panel that could be advantageous to introduce into current breeding programs. Further work investigating the extent of conditional neutrality relative to maladaptive pleiotropy in respect to local adaptation may be useful for future breeding efforts in perennial ryegrass, as this may inform breeders about the costs and benefits in selecting for increased local adaptation compared to broad adaptation.

## Acknowledgements

This study was carried out under the auspices of the ‘Public Private Partnership for pre-breeding in perennial ryegrass’ project (urn:nbn:se:norden:org:diva-12149), funded by the Nordic Council of Ministers, and the ‘Development of Molecular Markers for Genomic Selection of Adaptation in Perennial Ryegrass’ project funded by the Research Council of Lithuania (Grant No. MIP-64/2015). NordGen is acknowledged for their help obtaining the perennial ryegrass accessions for the project. NHJ was supported by a grant from the Aarhus University Research Foundation (Grant No. AUFF-F-2021-7-6).

---

**Supplementary Figure 1:**
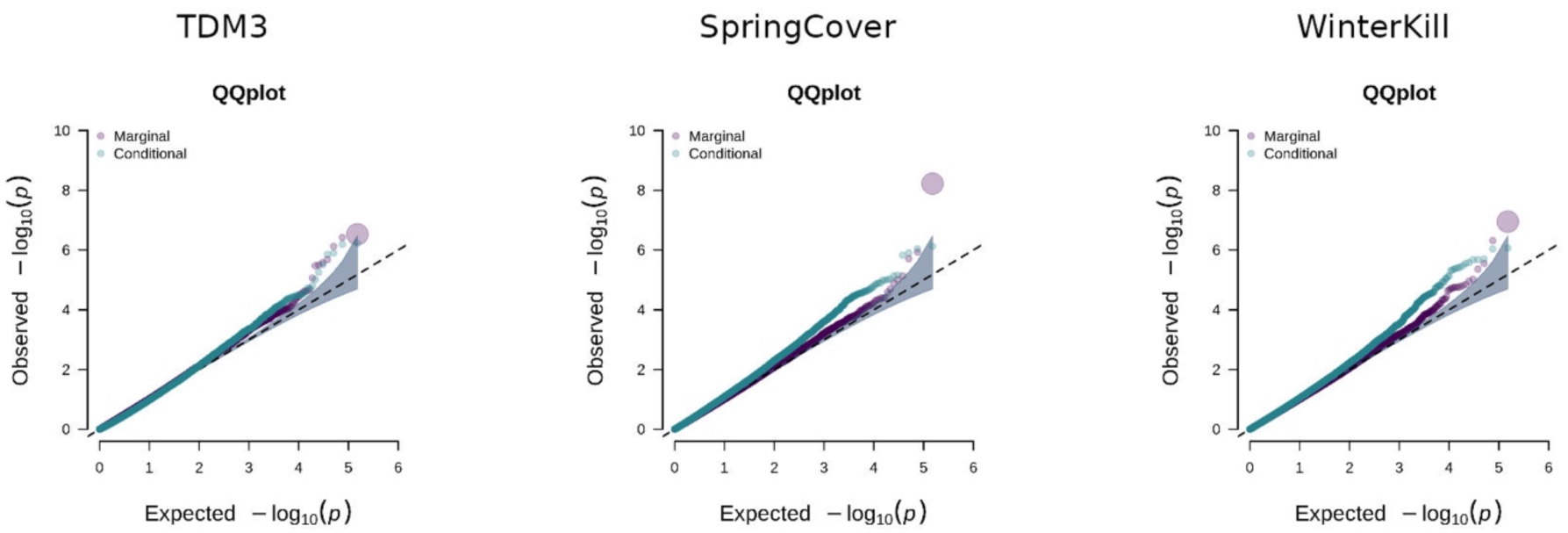
QQ-plots for the p-values from the GWAS analyses. P-values largely follow the diagonal, which suggests that our models effectively control for confounding factors like population stratification.

**Supplementary Table 1:**
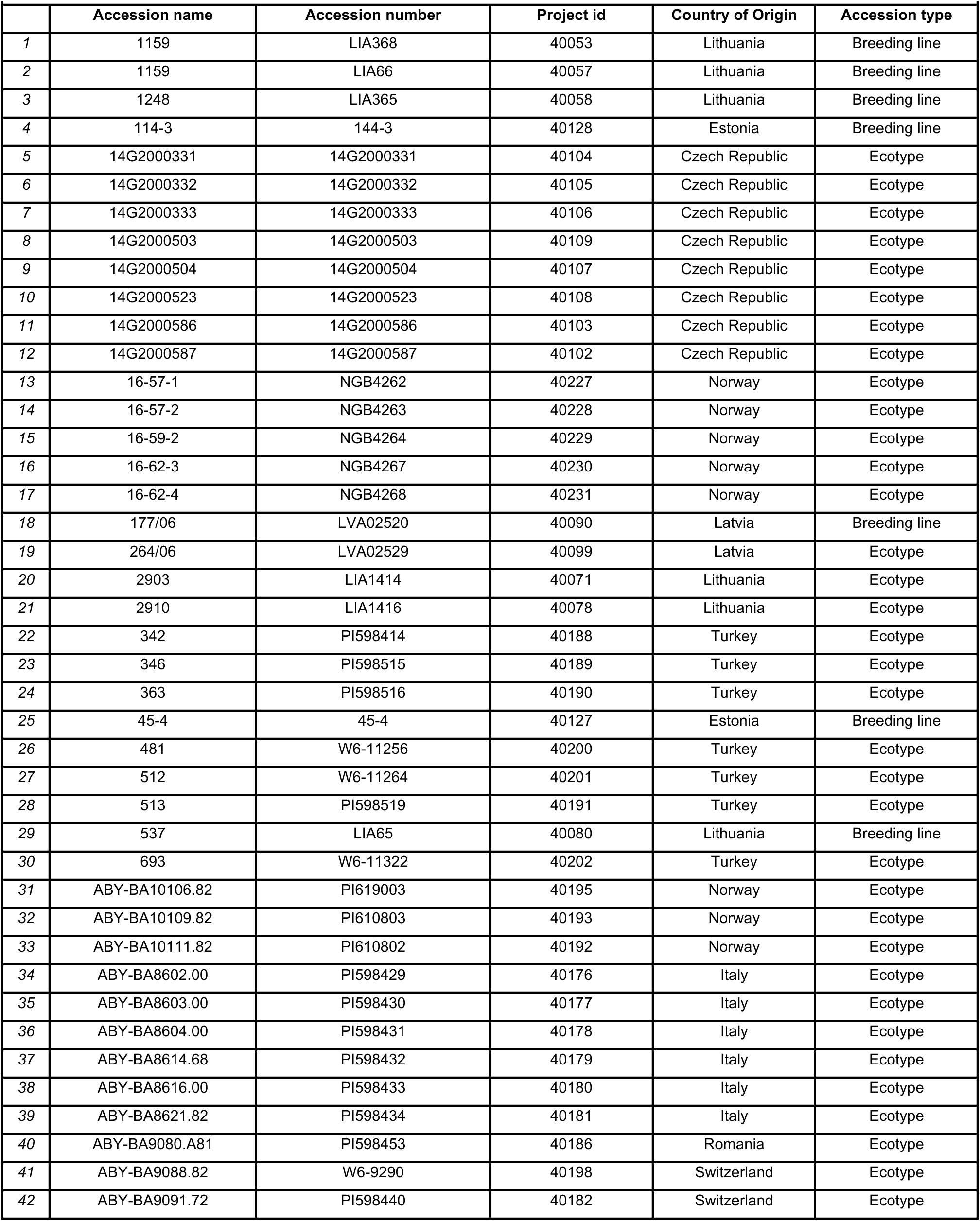
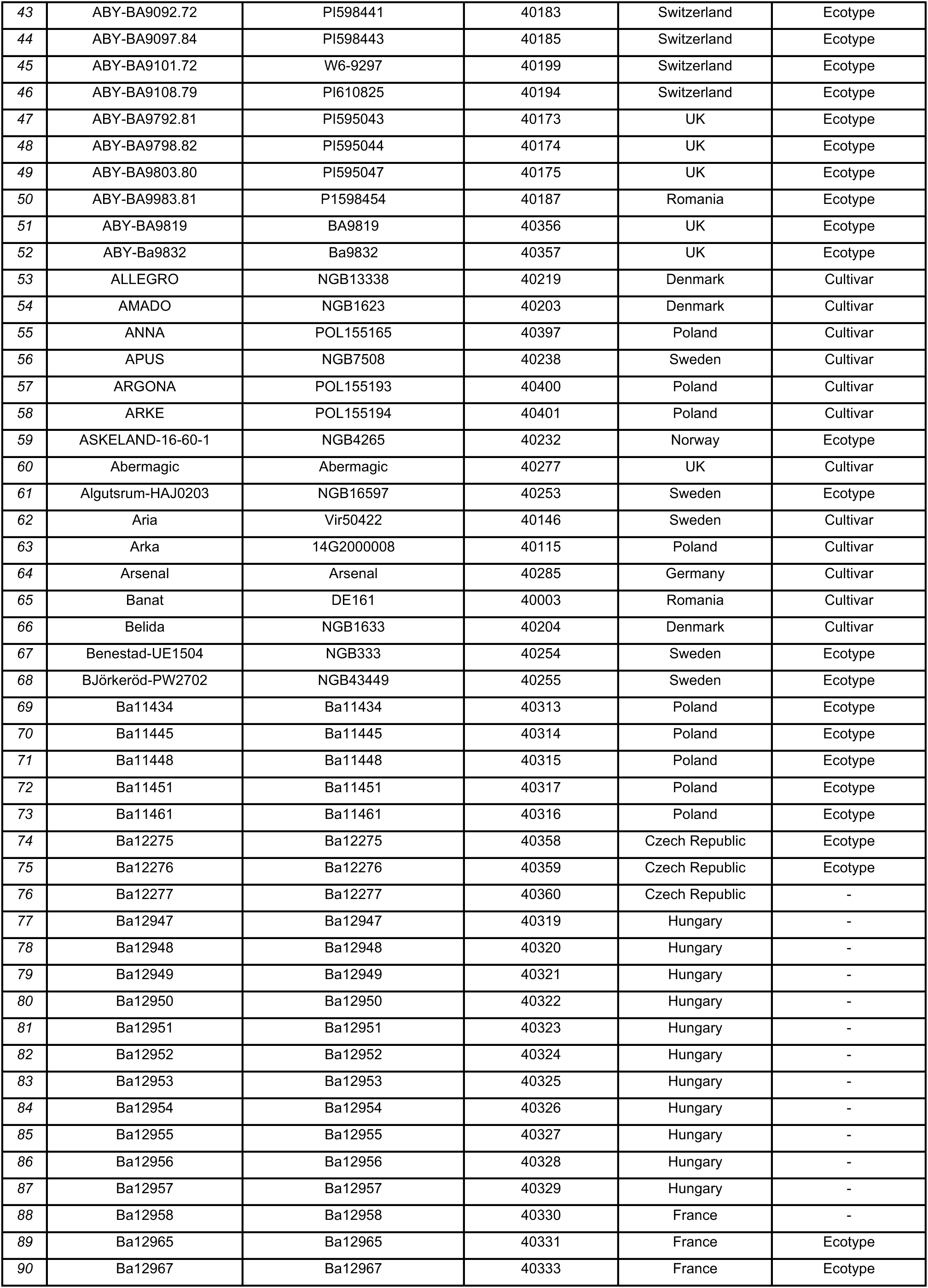
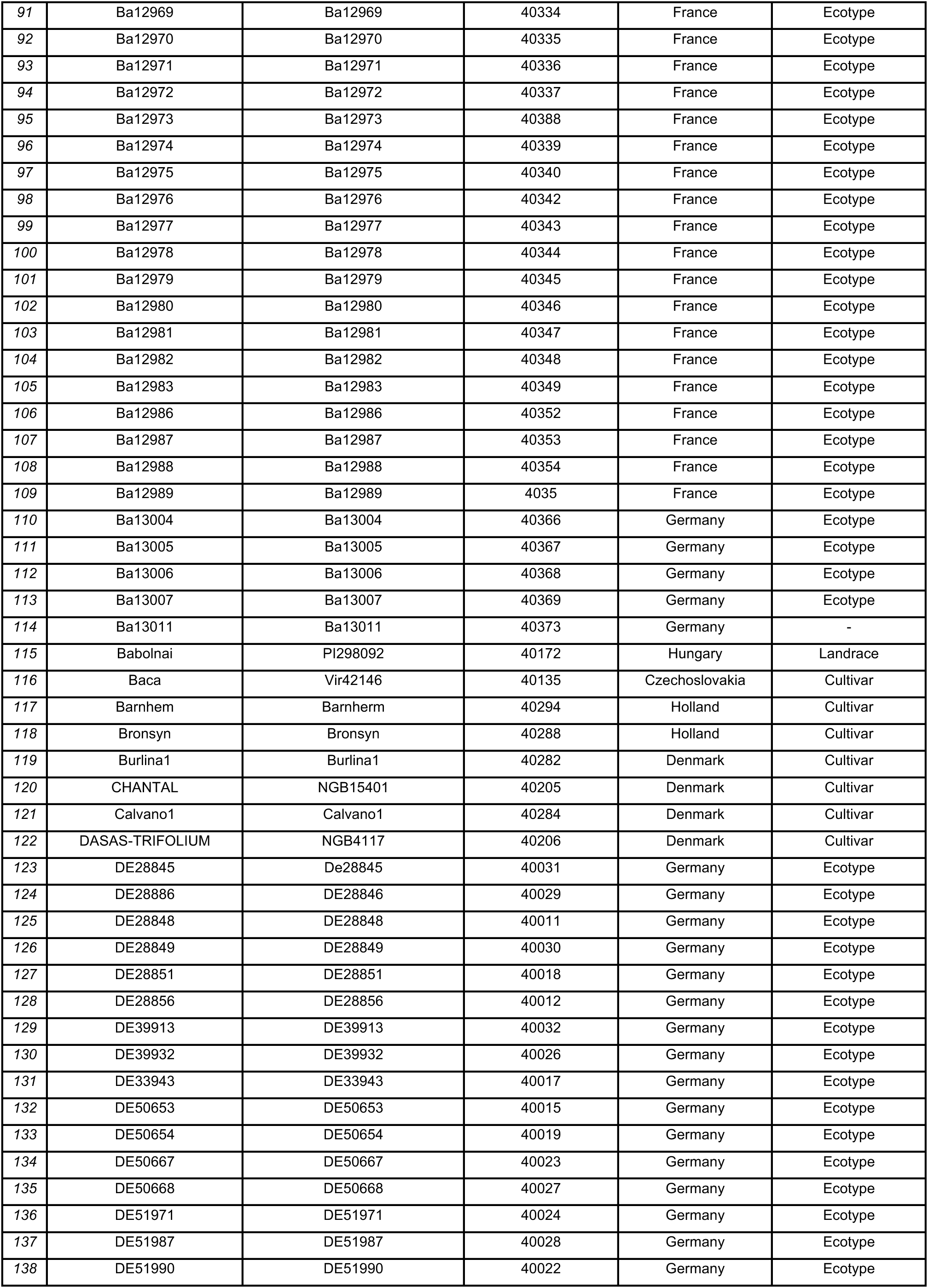
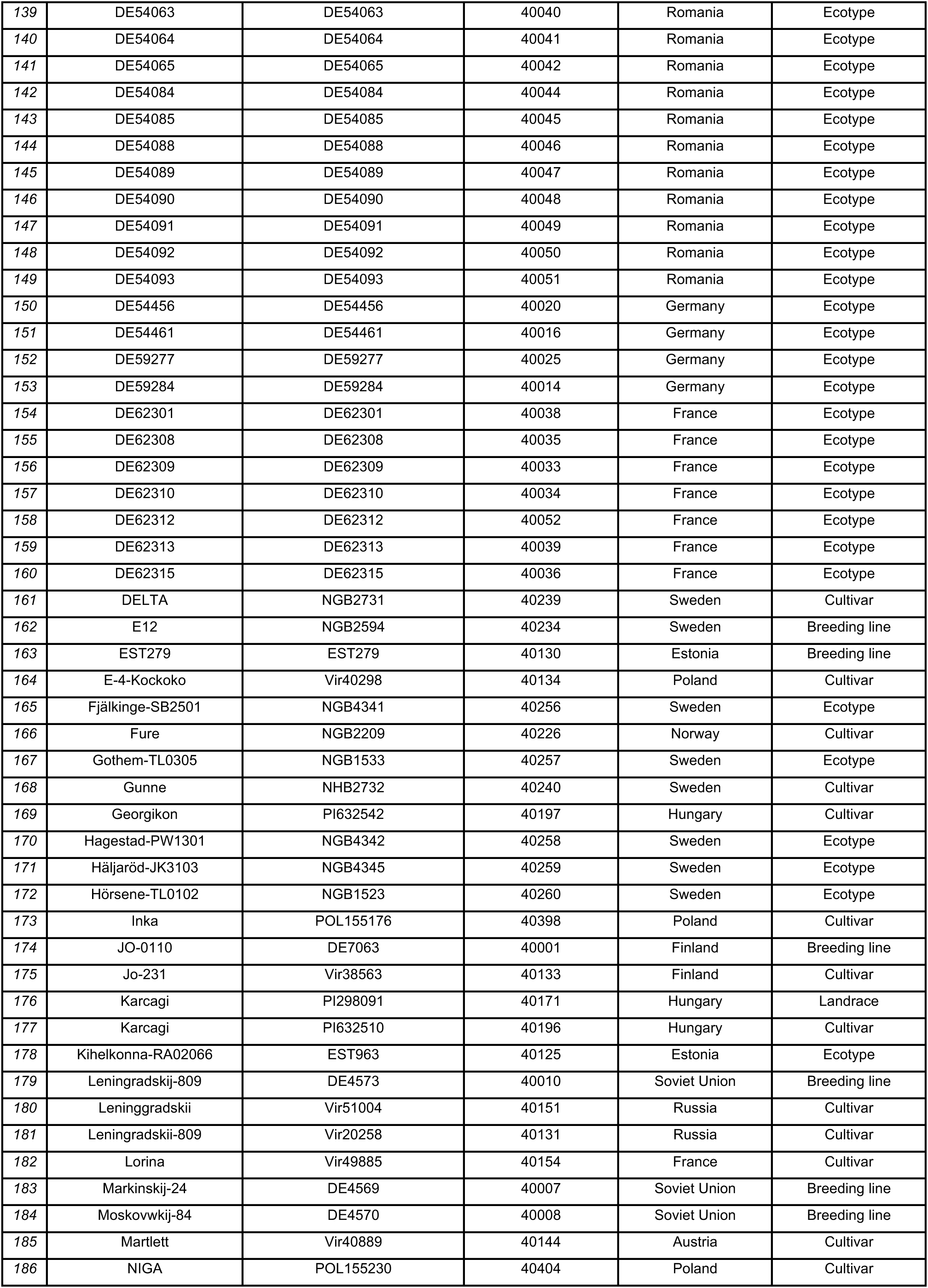
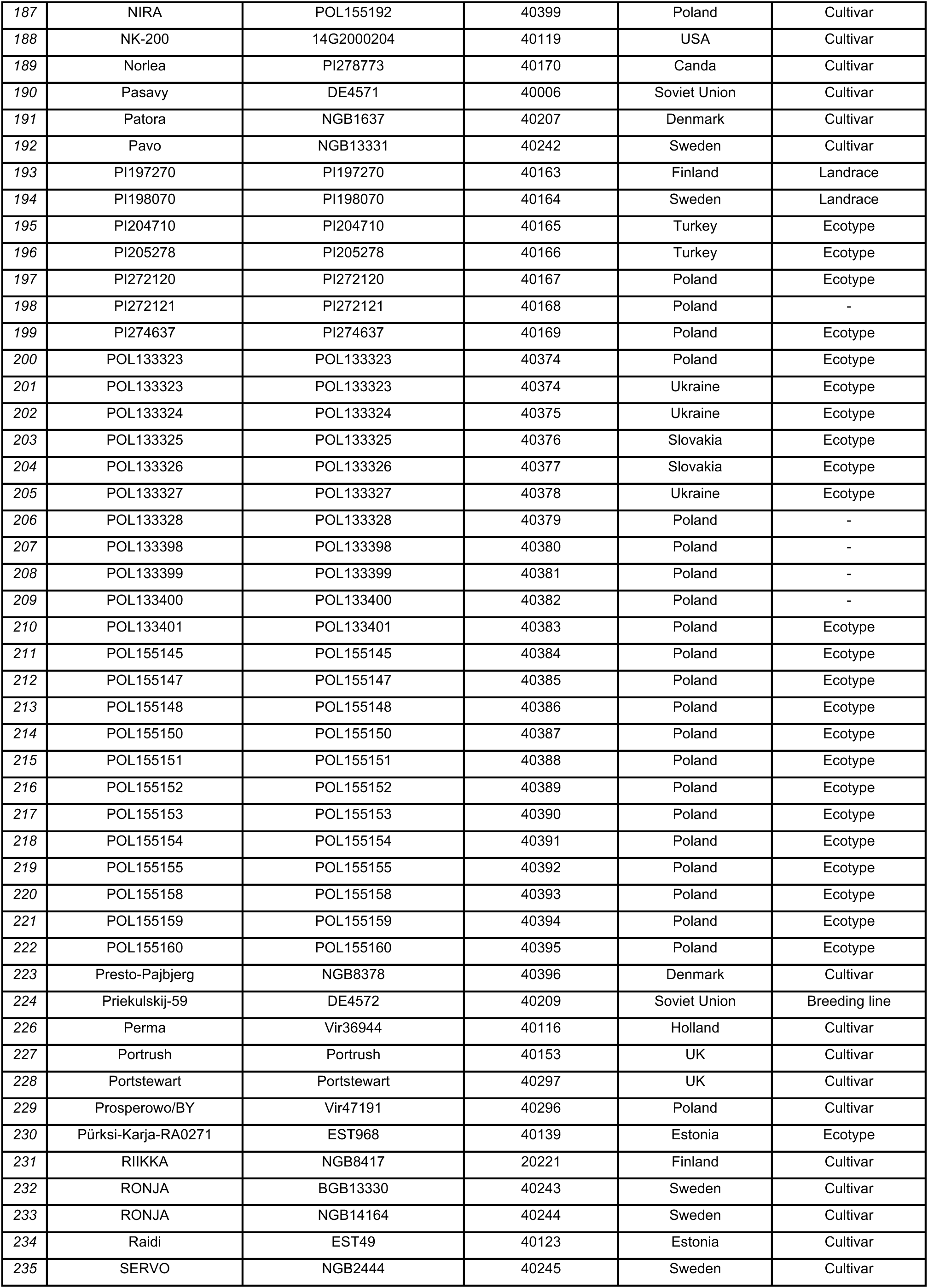
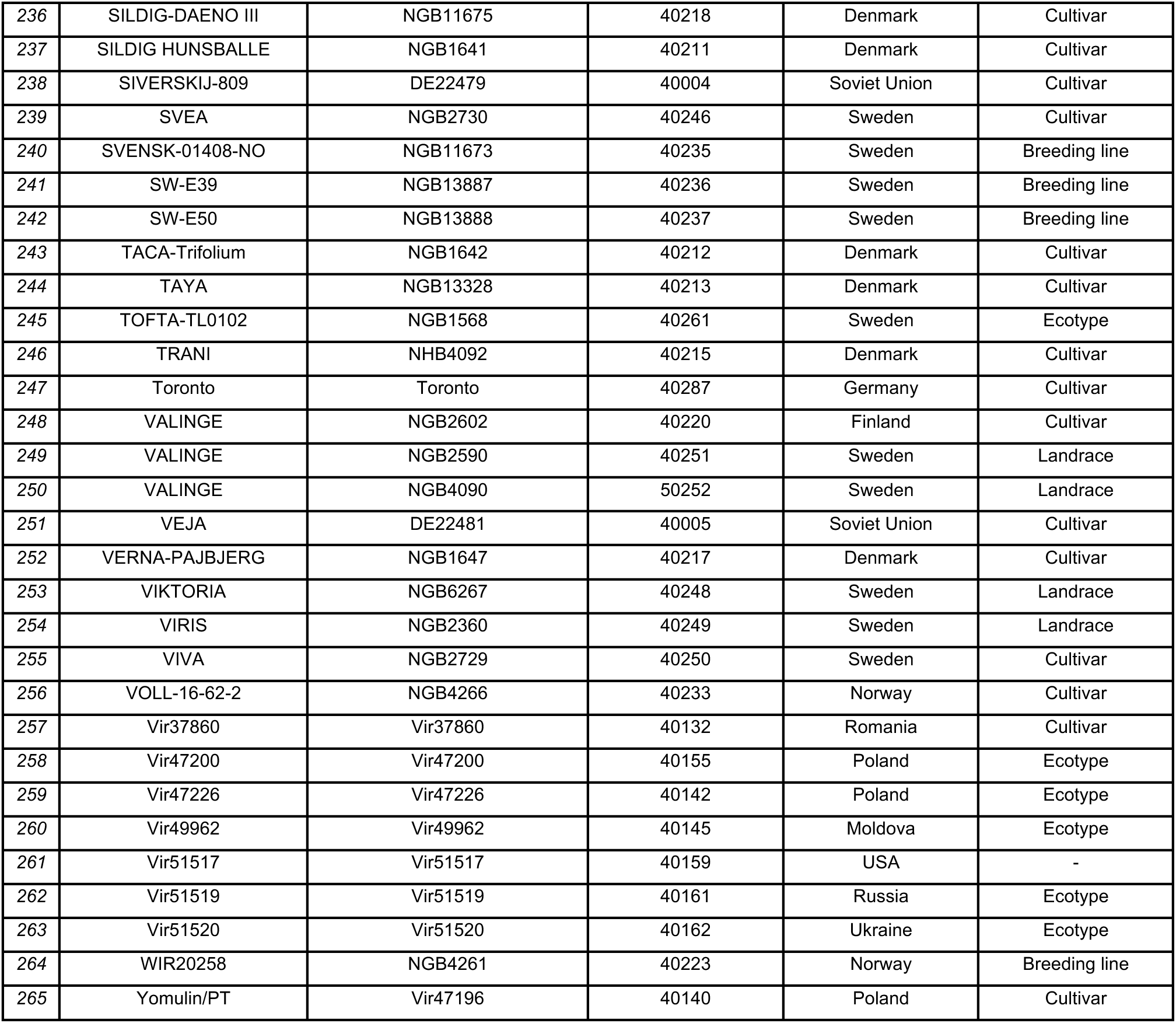
List of accessions names, project IDs, geographic origin and accession type (breeding line, landrace, cultivar, or ecotype) for the 265 diploid accessions included in the study.

**Supplementary Table 2:** List of environmental covariates used to construct the environmental similarity matrices used for the genomic prediction models.

| ID | Environmental covariate |
| --- | --- |
| ALLSKY_SFS_SW_DWN | All sky insolation incident on a horizontal surface |
| ALLSKY_SFC_LW_DWN | Thermal infrared longwave radiative flux |
| ETP | Evatransportation |
| FRUE | Effect of temperature and radiation use efficiency |
| n | Duration of sunshine hours |
| N | Daylight hours |
| PETP | Atmospheric water deficit |
| PRECTOT | Rainfall precipitation |
| RH2M | Relative air humidity at 2 Meters |
| RTA | Global Solar Radiation based on latitude and Julian day |
| SPV | Slope of saturation vapour pressure curve |
| T2MDEW | Dew-point temperature at 2 Meters |
| T2M | Temperature at 2 Meters |
| T2M_MIN | Minimum air temperature at 2 Meters |
| T2M_MAX | Maximum air temperature at 2 Meters |
| T2M_RANGE | Temperature range at 2 Meters |
| VPD | Vapour pressure deficiency |
| WS2M | Wind speed at 2 Meters |

**Table S3:** P-values for SNPs, below the genomewide threshold (FDR).

| Trait | Analysis | Chr | Position | P-value | FDR |
| --- | --- | --- | --- | --- | --- |
| SpringCover | Conditional | 1 | 137827458 | $1.49 \cdot 10^{-6}$ | 0.056 |
| SpringCover | Conditional | 2 | 312660071 | $7.54 \cdot 10^{-7}$ | 0.056 |
| SpringCover | Conditional | 4 | 106918014 | $1.25 \cdot 10^{-6}$ | 0.056 |
| SpringCover | Conditional | 4 | 291504000 | $9.14 \cdot 10^{-7}$ | 0.056 |
| SpringCover | Marginal | 2 | 340594974 | $5.99 \cdot 10^{-9}$ | 0.0009 |
| SpringCover | Marginal | 6 | 198771042 | $1.95 \cdot 10^{-6}$ | 0.098 |
| SpringCover | Marginal | 6 | 198771051 | $1.17 \cdot 10^{-6}$ | 0.088 |
| TDM3 | Conditional | 2 | 161529631 | $1.26 \cdot 10^{-6}$ | 0.052 |
| TDM3 | Conditional | 2 | 221602036 | $6.02 \cdot 10^{-7}$ | 0.048 |
| TDM3 | Conditional | 4 | 189930020 | $3.07 \cdot 10^{-6}$ | 0.092 |
| TDM3 | Conditional | 4 | 96059970 | $6.38 \cdot 10^{-7}$ | 0.048 |
| TDM3 | Conditional | 6 | 35366913 | $1.40 \cdot 10^{-6}$ | 0.052 |
| TDM3 | Marginal | 1 | 193676613 | $3.36 \cdot 10^{-6}$ | 0.072 |
| TDM3 | Marginal | 2 | 188828089 | $2.68 \cdot 10^{-6}$ | 0.072 |
| TDM3 | Marginal | 3 | 306736484 | $2.96 \cdot 10^{-7}$ | 0.028 |
| TDM3 | Marginal | 4 | 277056746 | $7.57 \cdot 10^{-7}$ | 0.038 |
| TDM3 | Marginal | 7 | 103894370 | $3.79 \cdot 10^{-7}$ | 0.028 |
| TDM3 | Marginal | 7 | 159935238 | $3.17 \cdot 10^{-6}$ | 0.072 |
| TDM3 | Marginal | 7 | 159935246 | $2.05 \cdot 10^{-6}$ | 0.072 |
| WinterKill | Conditional | 1 | 12416317 | $1.87 \cdot 10^{-5}$ | 0.098 |
| WinterKill | Conditional | 1 | 74714761 | $8.93 \cdot 10^{-6}$ | 0.075 |
| WinterKill | Conditional | 1 | 149556644 | $3.33 \cdot 10^{-6}$ | 0.052 |
| WinterKill | Conditional | 1 | 212715131 | $1.55 \cdot 10^{-6}$ | 0.094 |
| WinterKill | Conditional | 1 | 227248048 | $2.11 \cdot 10^{-6}$ | 0.052 |
| WinterKill | Conditional | 2 | 57085778 | $1.55 \cdot 10^{-5}$ | 0.094 |
| WinterKill | Conditional | 2 | 218558177 | $7.14 \cdot 10^{-6}$ | 0.072 |
| WinterKill | Conditional | 2 | 312660071 | $8.55 \cdot 10^{-7}$ | 0.052 |
| WinterKill | Conditional | 2 | 340594974 | $1.56 \cdot 10^{-5}$ | 0.094 |
| WinterKill | Conditional | 3 | 74496447 | $4.86 \cdot 10^{-6}$ | 0.052 |
| WinterKill | Conditional | 3 | 83644758 | $4.38 \cdot 10^{-6}$ | 0.052 |
| WinterKill | Conditional | 3 | 83644766 | $4.73 \cdot 10^{-6}$ | 0.052 |
| WinterKill | Conditional | 3 | 83644804 | $9.00 \cdot 10^{-6}$ | 0.075 |
| WinterKill | Conditional | 3 | 168639278 | $9.14 \cdot 10^{-7}$ | 0.052 |
| WinterKill | Conditional | 3 | 249015291 | $4.02 \cdot 10^{-6}$ | 0.052 |
| WinterKill | Conditional | 3 | 293400020 | $1.55 \cdot 10^{-5}$ | 0.094 |
| WinterKill | Conditional | 4 | 53655409 | $7.86 \cdot 10^{-6}$ | 0.074 |
| WinterKill | Conditional | 4 | 201753647 | $1.15 \cdot 10^{-5}$ | 0.085 |
| WinterKill | Conditional | 5 | 203021725 | $2.09 \cdot 10^{-6}$ | 0.052 |
| WinterKill | Conditional | 5 | 203021751 | $2.62 \cdot 10^{-6}$ | 0.052 |
| WinterKill | Conditional | 6 | 75411051 | $3.73 \cdot 10^{-6}$ | 0.052 |
| WinterKill | Conditional | 6 | 75411052 | $4.01 \cdot 10^{-6}$ | 0.052 |
| WinterKill | Conditional | 6 | 64101962 | $1.90 \cdot 10^{-5}$ | 0.098 |
| WinterKill | Conditional | 7 | 124140775 | $1.18 \cdot 10^{-5}$ | 0.085 |
| WinterKill | Conditional | 7 | 203027370 | $1.87 \cdot 10^{-5}$ | 0.098 |
| WinterKill | Conditional | 7 | 203277106 | $1.19 \cdot 10^{-5}$ | 0.085 |
| WinterKill | Conditional | 7 | 325317951 | $2.00 \cdot 10^{-6}$ | 0.052 |
| WinterKill | Marginal | 1 | 176887443 | $1.12 \cdot 10^{-7}$ | 0.016 |
| WinterKill | Marginal | 7 | 114853062 | $4.82 \cdot 10^{-7}$ | 0.036 |

